# TEMPO: Detecting Pathway-Specific Temporal Dysregulation of Gene Expression in Disease

**DOI:** 10.1101/651018

**Authors:** Christopher Michael Pietras, Faith Ocitti, Donna K. Slonim

## Abstract

While many transcriptional profiling experiments measure dynamic processes that change over time, few include enough time points to adequately capture temporal changes in expression. This is especially true for data from human subjects, for which relevant samples may be hard to obtain, and for developmental processes where dynamics are critically important. Although most expression data sets sample at a single time point, it is possible to use accompanying temporal information to create a virtual time series by combining data from different individuals.

We introduce TEMPO, a pathway-based outlier detection approach for finding pathways showing significant temporal changes in expression patterns from such combined data. We present findings from applications to existing microarray and RNA-seq data sets. TEMPO identifies temporal dysregulation of biologically relevant pathways in patients with autism spectrum disorders, Huntington’s disease, Alzheimer’s disease, and COPD. Its findings are distinct from those of standard temporal or gene set analysis methodologies.

Overall, our experiments demonstrate that there is enough signal to overcome the noise inherent in such virtual time series, and that a temporal pathway approach can identify new functional, temporal, or developmental processes associated with specific phenotypes.

**Availability:** An R package implementing this method and full results tables are available at bcb.cs.tufts.edu/tempo/.

## 1 Introduction

Understanding the dynamic aspects of molecular processes is essential, especially for inherently temporal functions such as those involved in development, disease progression, or aging (Przytycka et al., 2010; Yosef and Regev, 2011). Transcriptional profiling, whether by microarrays, RNA-seq, or other technologies, has proven useful for identifying temporal regulatory programs.

However, the csollection of data from large numbers of time points has proven to be prohibitively expensive and fraught, particularly in cases involving human subjects (Zinman et al., 2013). Thus the number of available data sets that include sufficient temporal resolution to solve key problems of interest remains limited. In most available human data sets, samples are taken only during medically indicated procedures, often yielding a single time point per individual.

If temporal information is available, however, it is possible to combine multiple samples from individuals at different ages or times into a single virtual time-series. Here we describe a method using temporal models of expression and functional gene sets to identify how and why those models break down in disease states. We do this using existing data sets featuring a single time point per individual, and we demonstrate that by so doing we can learn new things about the temporal and developmental processes associated with specific phenotypes.

### 1.1 Previous Work

The analysis of time series is a well-established field of data science whose relevance to expression data analysis has long been known. Computational methods specifically developed for the analysis of time series expression data are the subject of many papers and reviews (Spies and Ciaudo, 2015; Bar-Joseph, 2004; Bar-Joseph et al., 2012)). For example, several approaches to clustering temporal gene expression profiles have been proposed (e.g., (Ernst and Bar-Joseph, 2006; Androulakis et al., 2007; Bar-Joseph et al., 2012; Ramoni et al., 2002)).

Other methods have been designed to detect significantly different temporal expression profiles across experimental groups, conditions, or phenotypes. Most methods that do so (e.g., (Conesa et al., 2006; Bar-Joseph et al., 2003; Stegle et al., 2010)) use similar paradigms: each gene in each condition has an expression profile that is modeled as a function of time. A score is generated for each gene, capturing the difference between the models for the different conditions; genes are then ranked by their scores.

Most effective approaches, including those cited here, were designed specifically for time series expression data sets, which typically include only small numbers of samples for each condition and few time points. Notably, none of these methods explicitly scores gene *sets* or pathways, though it would be possible to adapt any of them to do so by using the gene scores as ranks and assessing gene set enrichment among the ranked gene lists.

However, of the methods we surveyed, only maSigPro (Conesa et al., 2006) provides publicly released code and has properties suitable for use with virtual time-series. Specifically, because virtual time series combine the availability of data from whatever time points appear in the static source data, they rarely feature matched case and control samples taken at consistent time points. This property rules out straightforward utilization of time series analysis methods that require the same set of time points across both conditions, that don’t allow for missing data, or that don’t allow for multiple samples at the same time point. How to adjust such methods or their input data to allow their use with virtual time series is not readily apparent.

### 1.2 Our Contributions

Here, we introduce an approach we call TEMPO (TEmporal Modeling of Pathway Outliers) to identify pathways or gene sets that show phenotype-associated temporal dysregulation. Given a gene expression data set where each sample is characterized by an age or time point as well as a phenotype (e.g. control or disease), and a collection of gene sets or pathways, TEMPO includes the following steps. First, for each set of genes in the gene set collection, it builds a partial least squares model to predict the age of the control samples as a function of the expression of the genes in that gene set. Prediction accuracy in controls is assessed by cross validation. It then uses the same model, trained on all the control samples, to predict age in the samples with the phenotype of interest. The gene sets are ranked by a scoring function that prioritizes models that predict age well in the controls but poorly in the disease samples, suggesting temporal dysregulation. We assess the significance of the observed scores via permutation.

Note that finding models that perform well in control samples but break down in other conditions is the underlying theme of several existing outlier detection methods, including our own (Noto et al., 2010, 2012). Such strategies have therefore been widely used in a variety of contexts. However, this is the first application of this methodology to temporal models of transcriptional profiles.

We compare the ranked lists of gene sets output by TEMPO to those from two other analyses of the same data sets: Gene Set Enrichment Analysis (GSEA) (Subramanian et al., 2005), a standard gene-set enrichment approach to differential expression analysis that makes no explicit use of temporal information, and maSigPro (Conesa et al., 2006), the only comparator method whose use on virtual time series data is straightforwardly feasible for the reasons indicated above. Still, because maSigPro itself does not look at functional enrichment, we need to translate its results at the gene level to the level of gene sets. To do so, we rank the genes by their maSigPro scores and then use GSEA to identify functional enrichment in the ranked list.

We demonstrate TEMPO’s utility on four previously published expression data sets, three of which examine peripheral blood in patients with neurological conditions. The first of these is a developmental microarray data set comparing gene expression in children with or without autism spectrum disorders. The next two data sets examine neurodegenerative disorders whose progression correlates with age: a microarray data set measuring expression in the blood of people with or without Alzheimer’s disease; and an RNA-seq data set that measures gene expression in adults of different ages with Huntington’s disease, either before or after the onset of symptoms, or in controls. The fourth data set looks at expression in airway epithelial cells of smokers with and without COPD.

We initially chose Gene Ontology (GO) Biological Process terms (Ashburner et al., 2000) as our gene set collection for the experiments described here. However, for the autism data set, we augmented the GO annotations with annotations from the DFLAT project, which incorporates additional developmentally relevant annotations into the GO framework (Wick et al., 2014).

Comparing the output of different analytical methods can be complex, because related functional terms often involve similar groups of genes, so the gene sets are not independent of each other. For example, if one method implicates “neuron apoptotic process” and another “regulation of neuron death,” two terms that share a common parent in the GO hierarchy (“neuron death,” GO:0070997), we would like to capture this relationship. We therefore use a measure based on semantic similiarity (Resnik, 1999) to assess relationships between the top gene-set lists output by different analytical methods.

Our examples demonstrate that TEMPO can identify age- and phenotype-related changes in expression that differ from those found by either the static analysis of GSEA or the traditional temporal modeling analysis in maSigPro. Further, our work illustrates the power of combining existing static data into virtual time series to study pathway-related temporal changes in dynamic processes.

## 2 Methods

### 2.1 TEMPO

#### 2.1.1 Computational model to predict age

For a gene set *G*, TEMPO trains a partial least squares regression (PLSR) model (Wold, 1985), using the *pls* package in R, to predict age as a function of the expression of all genes in *G*. Ages for all the control samples *C* = {*S*_1_, *S*_2_, *…S*_*j*_} are predicted in leave-one-out cross-validation using *j* separate PLSR models *M*_1_, *M*_2_, *…M*_*j*_ (Figure 1). PLSR models with up to 10 components were built for each gene set; we then chose the most accurate of these models in leave one out cross-validation on the control samples, and used that model for predicting ages in the test samples. (Note that this step is not illustrated in Figure 1 to improve readability.) The best *single* size is chosen and used to train one final model *M*_*j*+1_ on the control samples *C* = {*S*_1_, …, *S*_*j*_}. We then determine if the model is significantly predictive via permutation testing. For a gene set *G*, we compare to 500 randomly generated gene sets made up of |*G*| randomly selected genes, and train predictive models using those gene sets using the same process. We compare the control mean-squared error in cross validation for the model for *G* to that of each of the randomly generated gene sets, and say the model for *G* is significant if its control MSE is in the bottom 5% of the distribution of MSEs derived from these random sets (i.e., that the p-value associated with the control MSE for *G* is below 0.05). Then, if the model is significant by these criteria, ages for disease samples *D* = {*S*_*j*+1_, …, *S*_*k*_} are predicted using *M*_*j*+1_. Note that we also considered using other regression models in place of PLSR (see Appendix A), but we found PLSR to be most effective.

**Figure 1:**
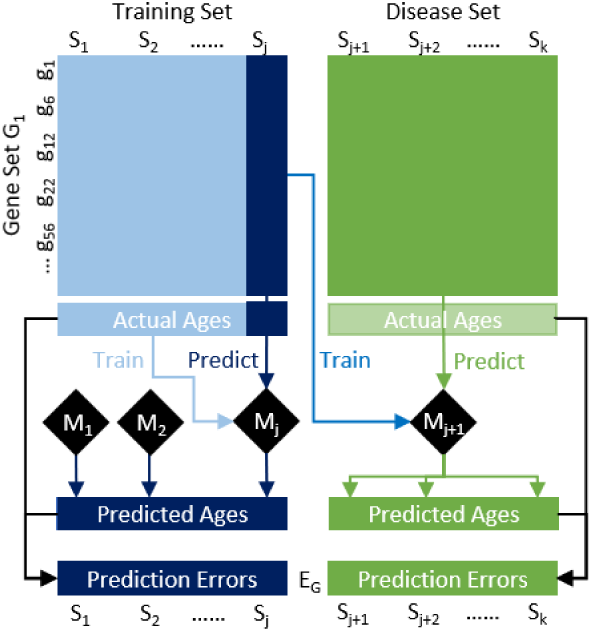
PLSR prediction for an arbitrary gene set *G*_1_. For *j* training samples, ages are predicted using *j* PLSR models in cross-validation. For the *k* − *j* disease samples, ages are predicted using a single PLSR model trained on all training samples. The difference between the predicted and actual ages for sample *S*_*i*_ is the prediction error 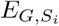.

#### 2.1.2 Scoring gene sets by performance on cases and controls

For each gene set *G* that has a significant model by the criteria described above, we have a set of age predictions for all control samples *C* and all disease samples *D*. We obtain a vector of prediction errors for *G*, the differences between the predicted ages for *G* and the actual ages. We call this vector of prediction errors *E*_*G*_, where *E*_*G,s*_ is the prediction error for sample *s* under gene set *G*. Using these errors, we determine the degree to which *G* is temporally dysregulated by calculating a score that incorporates the accuracy of the predictions for the control samples and the *in*accuracy of the predictions for the disease samples.

If our data sets behave as expected, these errors can be assumed to be normally distributed (although we assess and relax this assumption in Appendix B). Let *µ*_*G*_ and *σ*_*G*_ be the mean and standard deviation of the observed prediction errors on the control samples for gene set *G*, and let 𝒩_*G*_(*x*) be the probability of seeing an error at least as large as *x* under the normal distribution with mean *µ*_*G*_ and standard deviation *σ*_*G*_.

We then calculate the following score for gene set *G*, control sample set *C*, and disease sample set *D*:

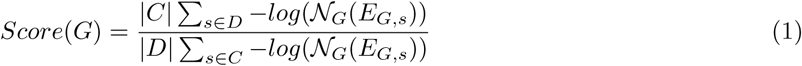

This is essentially a normalized ratio of the average “surprisal” score (Shannon, 1948) of the disease samples to that of the control samples. It is highest when the disease sample predictions are surprisingly bad, using an accurate model trained on the controls.

This score also captures our criteria for interesting gene sets. In gene sets where a reliable temporal pattern of expression in the controls breaks down in disease, we would be able to build a regression model that accurately predicts age in the control samples, but is unable to predict age accurately in disease, yielding many samples with improbable prediction errors and a high score. In gene sets where this is not the case, the regression model will have the same predictive power regardless of class label, yielding low scores (Figure 2).

**Figure 2:**
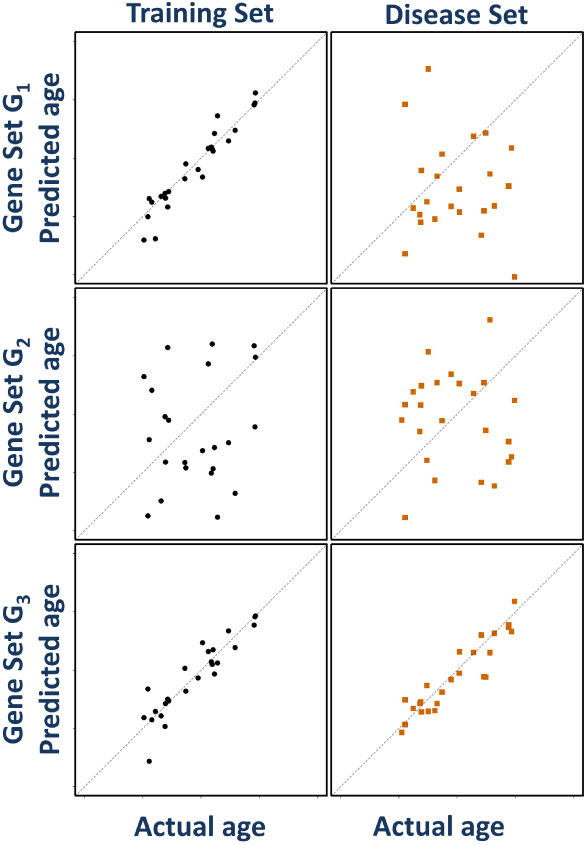
Predicted age v. actual age for hypothetical gene sets *G*_1_, *G*_2_, and *G*_3_ for control (left) and disease (right) samples. Gene sets like *G*_1_ have higher scores (Eq. 1).

The 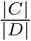 factor normalizes the score for the size of the control and disease sample sets, allowing meaningful comparison of results across experiments.

#### 2.1.3 Significance of Observed Scores

We estimate statistical significance via a permutation testing procedure. Specifically, we generate a set of 500 random permutations of size-matched gene sets. We permute by size-matched gene sets instead of the more traditional permutation of class labels because the ratio of the average surprisal scores used in our scoring function can be sensitive to differences in the age distribution between cases and controls, a common confounding factor in many data sets. Such differences can result in situations where even extremely poor models of age as a function of gene expression would be reported as significant, as in the hypothetical example in Figure 3.

**Figure 3:**
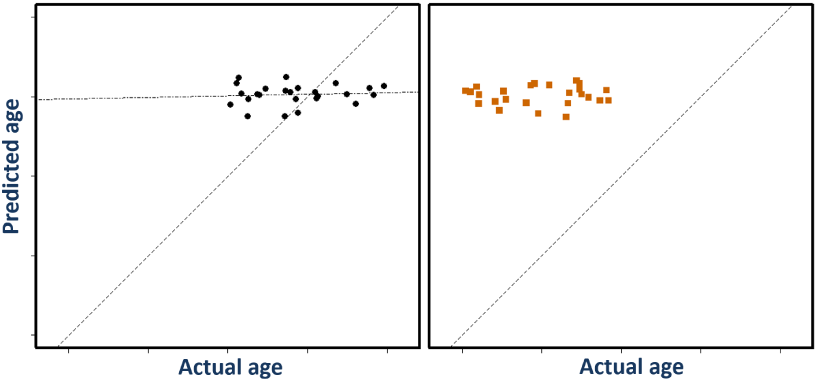
Predicted age v. actual age for a hypothetical gene set with no age-related signal, for control samples on the left and disease samples on the right. PLSR has no true predictive power in this gene set; it predicts almost the same age regardless of input. However, due to the different age distributions in the control and disease sets, the average surprisal ratio term of Equation 1 is relatively high, because the control predictions are close to the ideal *x* = *y* line, while the disease predictions are farther from it. Permuting by size-matched gene sets, rather than scrambling class labels, preserves this property in the permutations, ensuring that such gene sets are not inappropriately reported as significant.

For each permutation *P* in this set, we build a new temporal model on the same set of “control” samples and recompute the score of the gene set for that permutation (we call this *Score*(*P*)). The reported p-value for *G* is simply the percentage of all permutations where *Score*(*P*) ≥ *Score*(*G*). To account for multiple hypothesis testing, we calculate false discovery rates using the Benjamani-Hochberg procedure (Benjamini and Hochberg, 1995).

We report results for gene set *G* only if raw *p* ≤ 0.05 and FDR ≤ 0.25. Since this method is primarily intended for hypothesis generation, we might still be interested in gene sets with a false discovery rate this large; this is the default cutoff for the GSEA software as well (Subramanian et al., 2005). Both of these values (raw p and FDR) are reported in our full results tables online.

### 2.2 Expression data sets

#### Autism spectrum disorders

The autism data set, referred to as ASD, is based on a study by Mark Alter, *et al.* (Alter et al., 2011), that includes expression microarray data from peripheral blood lymphocytes for 59 control patients and 72 patients with autism spectrum disorders, with ages ranging from two to fourteen years. The data are available as GSE25507 in the Gene Expression Omnibus (GEO) database (Edgar et al., 2002); from this data set, we used all the samples for which subject ages were available.

#### Alzheimer’s Disease

The Alzheimer’s disease data set, referred to as AD, is based on a subset of the data used in a study by Sood, *et al.* (Sood et al., 2015) from the AddNeuroMed consortium (Lovestone et al., 2009). We include all samples from Batch 1 (available as GSE63060 on GEO) marked as “included in the case-control study,” for a data set consisting of blood gene expression data for 49 samples from Alzheimer’s patients and 67 from roughly similar-aged controls. All of these samples were annotated with patient ages in integer years.

#### Huntington’s disease

The Huntington’s disease data set, referred to as HD, includes normalized gene counts from an RNASeq experiment characterizing blood from Huntington’s disease patients (Mastrokolias et al., 2015). Its GEO accession number is GSE51779. The data set includes 33 control samples and 91 Huntington’s disease carriers, 27 of whom are asymptomatic (defined as patients for whom the motor score component of the Unified Huntington’s Disease Rating Scale (van Duijn et al., 2008) is 5 or less). All of these samples were annotated with patient ages in years to .01 precision, ranging from about 20 to 80 years.

#### COPD

The COPD data set is based on microarray data from studies by Carolan, *et al.* (Carolan et al., 2006) and Tilley, *et al.* (Tilley et al., 2009), available as GSE5058 on GEO. This data set contains small airway gene expression data from 15 smokers with COPD and 12 smokers who are apparently healthy. Each patient has an integer age in years.

### 2.3 Gene set collections

For the HD, AD, and COPD data sets, we used Gene Ontology (Ashburner et al., 2000) (GO) Biological Process gene sets. However, for the ASD data set, we used a version of the GO collection augmented with additional developmentally relevant annotations from the DFLAT project (Wick et al., 2014). Specifically, the Feburary 19, 2016 gene set gmt files were downloaded from the DFLAT web site (dflat.cs.tufts.edu). The Gene Ontology collection, generated at the same time as the DFLAT gene sets, was obtained from the same web site. Both the DFLAT and GO collections were filtered to remove all gene sets of size greater than 500 or less than 5, resulting in a total of 8416 DFLAT gene sets and 6484 GO gene sets.

### 2.4 Comparator methods

#### 2.4.1 GSEA

To account for differences in expression that are not related to age or time, we compare to Gene Set Enrichment Analysis (Subramanian et al., 2005). GSEA ranks gene sets by how represented genes from a given gene set are at the top (or bottom) of the list of all genes ranked by differential expression between two conditions. In this mode, using the actual expression data as input, GSEA does not account for any differences in expression as a function of time.

#### 2.4.2 maSigPro

To apply maSigPro to our temporal data sets, we first translated each of our static expression data sets into a suitable time series data set, with the number of replicates equal to the number of patients and each with a single time point.

We used the R package released with maSigPro (Conesa et al., 2006) to generate scores for each of the genes measured in each of our data sets. We then needed to extend these results to identify implicated *gene sets* rather than individual genes. We therefore used the “preranked” option in GSEA, with the rankings corresponding to the maSigPro scores, to identify differentially-expressed gene sets. It is worth noting that with preranked data, GSEA assesses significance by permuting gene sets, since it cannot permute class labels.

### 2.5 Comparing gene set lists

#### Semantic similarity

To compare the similarity of the top-scoring gene sets from different analyses, exact-match methods are insufficient, because different analyses may find different but related terms; one may discover “apoptotic process” while another may highlight “neuron apoptotic process.” To capture these semantic relationships, we use pairwise Resnik semantic similarity scores (Resnik, 1999). All scores were calculated using the GoSemSim (Yu et al., 2010) R package. Although GoSemSim offers tools for calculating semantic similarity between sets of GO terms, we found these numbers difficult to assess in absolute terms.

To address this, we instead examine which terms have significantly similar matches in the other term set. That is, given two collections of terms *T*_1_ and *T*_2_, for each term *t*_*i*_ in *T*_1_, we want to know if there exists a semantically similar term *t*_*j*_ in *T*_2_. Given the distribution of pairwise Resnik similarity scores involving *t*_*i*_, we say a term *t*_*j*_ is semantically similar to *t*_*i*_ if Resnik(*t*_*i*_,*t*_*j*_) is above some chosen cutoff *c*. For our experiments here, we chose *c* = 0.6, which corresponds to approximately the top 0.3% of pairwise Resnik scores between all biological process gene sets, and compare between collections of gene sets of size 40. The number 40 was not tuned, but was chosen (somewhat arbitrarily) to represent a good variety of top functions in the output.

We note that it is likely that some number of gene set pairs between two collections are semantically similar by chance alone. We quantify this likelihood by permutation. For each permutation *i*, we generate two random collections of 40 terms and determine the number of terms *n*_*i*_ from the first collection that have semantically similar terms in the second. We do this 500 times, and compare the number of similar terms *n* from *T*_1_ to *T*_2_ to this distribution to obtain the likelihood of seeing as much similarity by chance; this is simply the fraction of permutations where *n*_*i*_≥ *n*. For |*T*_1_| = |*T*_2_| = 40, we found that an overlap of at least 8 semantically similar gene sets is required for the likelihood of seeing such overlap by chance to be below 0.05.

#### Correlation

We also consider the Spearman’s rank-correlations between full gene set lists from two different analyses. While such an approach penalizes changes in the rankings of even insignificant gene sets, it has the advantage that it involves all gene sets equally. While high TEMPO and high GSEA scores denote something comparable, low TEMPO scores denote a lack of temporal expression patterns and low GSEA scores can indicate enrichment in the control condition. Thus we do not consider Spearman correlations between either GSEA or maSigPro and TEMPO to be meaningful. Using the absolute value of the GSEA score might be more appropriate for such comparisons, but again it is not clear that such values would be comparable with rankings by other methods.

## 3 Results and Discussion

### 3.1 TEMPO finds unique temporal dysregulation in disease classes

In all four data sets, TEMPO identifies pathways that are known to change with age, but whose normal temporal trajectory is disrupted in disease. The observed temporal dysregulation is in many cases consistent with prior knowledge and sometimes consistent with identified or proposed therapeutic targets for treating the indicated disease. Thus, novel findings from this approach may suggest possible new targets or interventions.

The TEMPO results differ in many respects from the gene sets returned by comparator methods GSEA and maSigPro. *No exactly identical* gene sets appeared in the top 40 listed in any TEMPO analysis and any comparator method. Table 1 shows the number and significance of *semantically similar* gene sets observed between TEMPO and the comparator methods. Furthermore, in several cases, either GSEA or maSigPro does not identify *any* significant gene sets. In such cases, we nonetheless compared the semantic similarity of the top 40 highest-scoring gene sets from each method to those in the TEMPO results.

**Table 1:** Number of semantically similar gene sets (with significance) from the top 40 TEMPO results for the data set in each row and the top 40 results of the indicated comparator method run on the same data set.

| TEMPO on | GSEA | maSigPro+GSEA |
| --- | --- | --- |
| Autism | 0 (1.000) | 1 (0.958) |
| Huntington’s | 0 (1.000) | 1 (0.958) |
| Alzheimer’s | 0 (1.000) | 0 (1.000) |
| COPD | 2 (0.838) | 1 (0.958) |

The differences between the TEMPO and GSEA results are not unexpected. Gene sets with high GSEA scores will not necessarily have high TEMPO scores, because gene sets where there is no pattern of expression as a function of time will not be scored highly by TEMPO regardless of any time-independent differential expression that may exist.

For space reasons, full results tables and scatter plots for all methods and data sets are available online at bcb.cs.tufts.edu/tempo/tempoV4/ and top results for all methods and data sets are available in the supplemental material. However, we discuss some example results for each data set, and we reproduce part of the TEMPO results table for the ASD data set in the main manuscript as an example.

### 3.2 ASD developmental dysregulation: neurotransmitters and inflammation

In the ASD expression data, TEMPO identified 235 significant gene sets. A selection of the highest-scoring of these is shown in Table 2. Common themes in this list include inflammation, angiogenesis, *PTEN* activity, developmental processes, and neurotransmitter signaling.

**Table 2:** A selection of high-scoring gene sets in ASD, ranked by TEMPO score.

| Rank | Gene Set | Control<br>MSE | Score | MSE<br>p-value | Score<br>p-value | Score<br>FDR |
| --- | --- | --- | --- | --- | --- | --- |
| 1 | positive regulation of glomerulus development | 3.816 | 3.899 | 0.036 | 0.002 | 0.075 |
| 2 | secretion by cell | 2.760 | 2.581 | 0.004 | 0.004 | 0.075 |
| 4 | phosphatidylinositol biosynthetic process | 3.224 | 2.456 | 0.004 | 0.004 | 0.075 |
| 5 | positive regulation of growth | 3.076 | 2.405 | 0.008 | 0.002 | 0.075 |
| 8 | phospholipid metabolic process | 3.154 | 2.343 | 0.012 | 0.002 | 0.075 |
| 9 | myeloid leukocyte activation | 3.092 | 2.343 | 0.008 | 0.002 | 0.075 |
| 11 | phosphatidylinositol metabolic process | 3.522 | 2.310 | 0.036 | 0.004 | 0.075 |
| 13 | positive regulation of sequence-specific DNA<br>binding transcription factor activity | 2.746 | 2.283 | 0.006 | 0.006 | 0.075 |
| 14 | regulation of integrin-mediated signaling pathway | 2.932 | 2.274 | 0.002 | 0.004 | 0.075 |
| 19 | inflammatory response | 3.281 | 2.189 | 0.018 | 0.010 | 0.086 |
| 22 | negative regulation of neurotransmitter uptake | 3.248 | 2.169 | 0.002 | 0.002 | 0.075 |
| 23 | regulation of serotonin uptake | 3.248 | 2.169 | 0.004 | 0.002 | 0.075 |
| 24 | negative regulation of serotonin uptake | 3.248 | 2.169 | 0.004 | 0.002 | 0.075 |
| 27 | myeloid dendritic cell activation | 3.036 | 2.139 | 0.006 | 0.006 | 0.075 |
| 31 | central nervous system neuron differentiation | 3.333 | 2.122 | 0.020 | 0.012 | 0.086 |
| 32 | phospholipid biosynthetic process | 3.324 | 2.120 | 0.016 | 0.010 | 0.086 |
| 40 | nervous system development | 3.225 | 2.090 | 0.048 | 0.010 | 0.086 |
| 44 | positive regulation of extrinsic apoptotic<br>signaling pathway in absence of ligand | 3.421 | 2.046 | 0.006 | 0.004 | 0.075 |
| 46 | cytokine production | 3.470 | 2.028 | 0.018 | 0.010 | 0.086 |
| 48 | positive regulation of cysteine-type endopeptidase<br>activity involved in apoptotic process | 3.418 | 2.021 | 0.024 | 0.014 | 0.086 |

Results from both a static GSEA analysis and the maSigPro-plus-enrichment analysis on the same data set are also available on the TEMPO web site. Neither GSEA nor maSigPro analysis returns *any* gene sets with FDR ≤ .25, though both have several hundred gene sets with raw *p* ≤ .05.

The role of serotonin and other neurotransmitters in the etiology of ASD has long been investigated (Ritvo et al., 1970; Cook et al., 1997). Although serotonin activity is evident very early in human development (Murrin et al., 2007), the nature and expression of serotonin response pathways change considerably during both childhood and adolescence (Crews et al., 2007), consistent with observations that children and adults respond differently to drugs targeting this system (Varigonda et al., 2015). Further, while SSRIs are often used to treat ASD patients, there is considerable evidence of increased adverse events in the pediatric autistic population, suggesting increased care is needed in the use of these drugs (Kolevzon et al., 2006). Understanding specifically how the expression of serotonin-related genes is expected to change with age in the neurotypical population, and how autistic patients differ from these expectations, may be key to the better prediction of tolerance and appropriate dosage in this population.

Figure 4 plots the actual and TEMPO-predicted ages for the gene set “regulation of serotonin uptake.” The plot on the left shows the relatively accurate predictive age models in the controls, while that on the right show how the developmental program of the genes in the pathway breaks down in the group of subjects with ASD.

**Figure 4:**
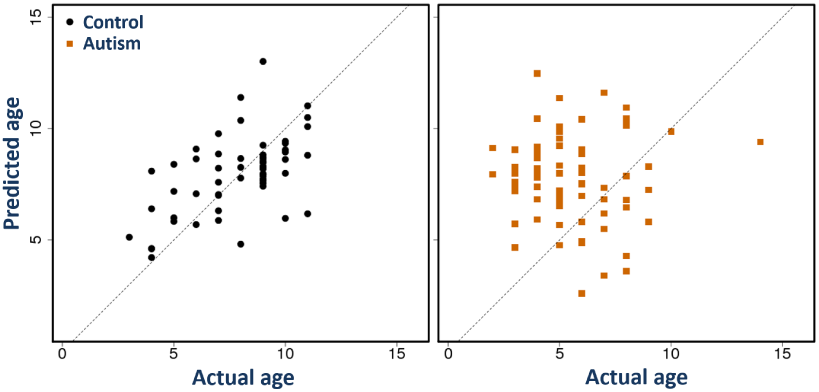
Predicted age vs. actual age for both control (black circles, left) and autistic (orange squares, right) subjects for genes with the annotation “Regulation of serotonin uptake” on the ASD data. Each dot represents one patient. Predicted ages are those produced by TEMPO using the model built from the controls, based only on the expression values of genes in the regulation of serotonin uptake pathway.

Inflammatory pathways have also been linked to ASD (Croonenberghs et al., 2002), and an increase in the circulating frequency of myeloid dendritic cells, which modulate immune response, has been observed in children with ASD compared to controls (Breece et al., 2013). NFKB signaling has been implicated as well (Ziats and Rennert, 2011), possibly contributing to the dysregulation of inflammatory cytokines (Lawrence, 2009).

Programmed cell death is known to play a key role in normal brain development (Yeo and Gautier, 2004). Disruption of apoptotic pathways has been shown to contribute to the development of ASD and to symptoms suggestive of it in animal models (Margolis et al., 1994; Wei et al., 2014). It has been suggested that abnormal *PTEN* function, which has been documented in a subset of the autism patients (Kyrylenko et al., 1999), may contribute to apoptosis in neural development by regulating PI3K / AKT signaling (Zhou and Parada, 2012; Wei et al., 2014). Both *PTEN* (also known as “phosphatase and tensin homolog”) and *PI3K* (“phosphatidylinositol-3-kinase”) are involved in phosphatidylinositol metabolism; this pathway has even been suggested as a possible therapeutic target for autism (Enriquez-Barreto and Morales, 2016). *PTEN* has also been shown to regulate angiogenesis (Choorapoikayil et al., 2013), which has itself been implicated in autism spectrum disorders (Azmitia et al., 2016).

### 3.3 Temporal dysregulation of apoptosis in Alzheimer’s disease

In the Alzheimer’s data set, TEMPO identified 140 significant gene sets. The pathways implicated include several processes known to have relevance in Alzheimer’s disease, including apoptosis, immunity, the DNA damage response, and regulation of phosphorylation.

Amyloid beta plaques have been observed to induce apoptosis in Alzheimer’s disease (AD) patients (Ghavami et al., 2014). Intrinsic apoptosis through altered mitochondrial permeability, triggered by accumulations of amyloid beta precursor protein, has been proposed as the mechanism by which amyloid plaques induce mitochondrial oxidative stress in AD Bartley et al. (2012). Four of the top 40 and 16 of the 140 significant gene sets identified by TEMPO in the Alzheimer’s population are related to apoptosis, including “positive regulation of apoptotic signaling pathway,” “regulation of apoptotic signaling pathway,” “intrinsic apoptotic signaling pathway,” and “regulation of intrinsic apoptotic signaling pathway,” with scores ranking 8th, 9th, 17th, and 25th, and all with FDR ≤ 0.1.

The substantial role of the immune system and cytokine signaling in Alzheimer’s is well explored (Rubio-Perez and Morillas-Ruiz, 2012), and has been proposed as the basis of new immunotherapeutic approaches (Monsonego et al., 2013). Previous work has shown changes in immune processes and signaling in healthy aging. For example, T-cell populations change and pro-inflammatory cytokine signaling increases with age (Garg et al., 2014; van der Geest et al., 2014). TEMPO’s identification of cytokine signaling and T-cell activation pathways in this context confirms that it is finding likely pathways that have a predictable age-related pattern that breaks down in disease, and that may suggest therapeutic targets.

### 3.4 Age-related expression dysregulation in pre-symptomatic HD patients

Huntington’s disease (HD) is known to be caused by a trinucleotide repeat expansion of the *huntingtin* (*HTT*) gene. However, many other genes have been found to modify the effects of these expansions, reflecting age of onset, severity, and specific characteristics of the disorder (Munoz-Sanjuan and Bates, 2011). Such modifiers are actively sought as potential avenues for devising new treatment approaches.

For this data set, 166 gene sets met the significance criteria, suggesting that there are age-specific expression patterns for many biological processes that are disrupted in the disorder. In contrast, neither Gene Set Enrichment Analysis nor maSigPro returns any significant gene sets for this data set. The full results for all these analyses are available at the TEMPO web site.

Two of the ten highest scoring gene sets in the TEMPO analysis are “regulation of ERBB signaling pathway” and “regulation of EGFR signaling pathway.” Prior evidence has implicated the ERBB pathway and EGFR signaling in the pathogenesis of HD (Kalathur et al., 2012; Liu YF, 1997). Mechanistic studies suggest that mutant *HTT* interferes with EGFR signaling, and ERBB signaling defects have been implicated in other neurodegenerative diseases including Alzheimer’s (Bublil and Yarden, 2007). Ion channel signaling, another top-scoring function, is known to be affected in HD (Wong et al., 2008), but whether this is a cause of or a reaction to Huntington’s pathology is not yet known(Mackay et al., 2018).

Another notable observation is the relative temporal dysregulation of telomere maintenance genes in HD, consistent with the second-ranked TEMPO hit “telomere maintenance via recombination.” Telomere length in HD has recently been verified to be shorter than in controls, and more so than in other forms of dementia (Kota et al., 2015). This process is known to reflect aging in general, but identifying further disruption of the normal aging patterns in HD represents an important finding with potential therapeutic implications.

These results are based on comparing controls to both symptomatic and pre-symptomatic patients together, but many of the same observations hold when symptomatic and pre-symptomatic patients are considered separately. The pairwise Spearman correlations between the TEMPO scores for just symptomatic, just pre-symptomatic, and the combined data set are all extremely high (≥ 0.99). The top-scoring gene sets returned by TEMPO for each of these three comparisons are also very similar, with a minimum of 21 out of the top 40 gene sets being semantically similar or identical in each pairing, as shown in Table 3. In general, there is more significant disruption of age-specific patterns in the symptomatic patients, but such disruptions are still detectable when comparing the pre-symptomatic patients to the controls (see e.g. Figure 5).

**Table 3:**
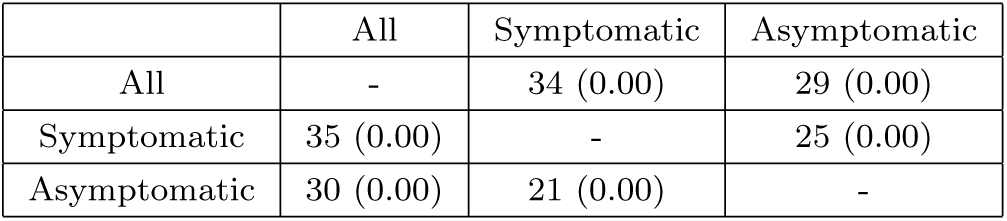
Number of semantically similar or identical gene sets (with significance) from the top 40 TEMPO results for the Huntington’s subset in each row to the top 40 TEMPO results for the subset in the corresponding column.

|  | All | Symptomatic | Asymptomatic |
| --- | --- | --- | --- |
| All | - | 34 (0.00) | 29 (0.00) |
| Symptomatic | 35 (0.00) | - | 25 (0.00) |
| Asymptomatic | 30 (0.00) | 21 (0.00) | - |

**Figure 5:**
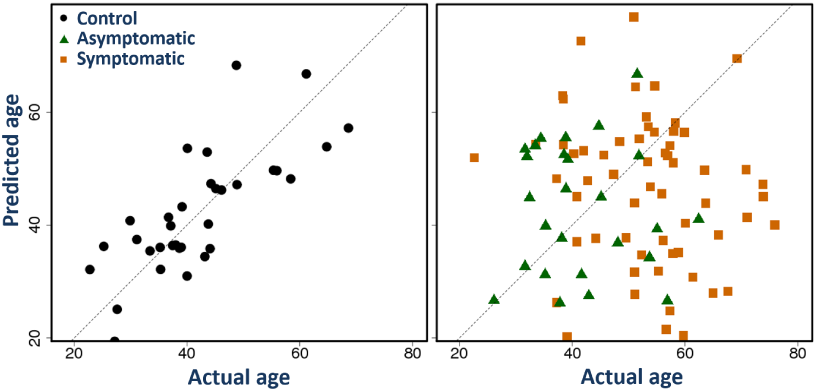
Predicted age vs. actual age for control (left), pre-symptomatic, and symptomatic Huntington’s (right) subjects for a top-scoring pathway on the Huntington’s Disease data, “Regulation of ERBB signaling pathway.” Each dot represents one patient in the control group; squares represent HD patients, and triangles represent HD patients who are still pre-symptomatic. Predicted ages are those produced by TEMPO using the model built from the controls, based only on the expression values of genes in the EERB signaling pathway.

Our results suggest a pattern of expression disruption for many of these gene sets that is detectable before disease onset. This is perhaps not surprising; prior imaging work has identified differential aging in a transgenic rat model of HD (Blockx et al., 2011), even before the onset of symptoms, and some critical expression changes have been documented in pre-symptomatic human HD patients (Chang et al., 2012). Still, the presence of a coherent change in age-related regulation prior to symptom onset may yield novel therapeutic insights.

### 3.5 Age-dependent dysregulation of immune pathways in COPD

In the COPD data set, TEMPO identified 176 significant gene sets whose predictable age-related expression relationships in airway epithelial cells from healthy smokers are disrupted in COPD.

Previous work has shown increased expression of pro-inflammatory cytokines and decreased NK cell activity in asymptomatic smokers (Zeidel et al., 2002). Increased inflammatory signaling correlates with pack-years, and therefore with age, even in smokers without apparent disease (Hacievliyagil et al., 2013). Unexpectedly early aging-like changes in vascular smooth muscle cells have been correlated with the inflammatory cytokines and oxidative stress likely to result from smoking (Trindade et al., 2017).

Consistent with this prior information, TEMPO finds excellent predictive models of age using the GO gene sets “positive regulation of interferon-gamma secretion,” “T-helper cell lineage commitment,” and “vascular smooth muscle cell development” that break down in COPD. These gene sets are ranked 2, 4, and 7 of those identified in the COPD data (ranking of gene set G is by Score(G)); the raw p-values for all of these are 0.004, and the corresponding adjusted FDR is less than 0.01.

Age related changes specific to alanine and glutamine transport have been observed in rat blood cells (Felipe et al., 1992). Although there is little data describing amino acid transport with age in human airway epithelial cells, it is intriguing that exactly these two have been observed with disrupted age related patterns in the COPD patients. Other immune and inflammatory processes, including activin receptor signaling, regulation of adaptive immunity, cytokine signaling, and B-cell activation, were found to be significantly disrupted in the COPD data as well.

### 3.6 Modest similarities between Huntington’s and Alzheimer’s

The number of semantically similar gene sets between the top TEMPO results in each of the three data sets from peripheral blood in neurodevelopmental or neurodegenerative disorders is not significant, with the maximal overlap between Alzheimer’s and Huntington’s, which share just two *identical* gene sets within the top 40 (“ammonium ion metabolic process” and “positive regulation of transporter activity”). However, the TEMPO scores between these two data sets are modestly Spearman rank correlated (0.309), and the control mean-squared errors for the age models in these two data sets are also modestly rank correlated (0.307). No other pair of data sets has as strong a similarity between either the TEMPO scores or the control mean-squared errors. This correlation is reasonable because the Alzheimer’s and Huntington’s disease data sets both feature older patients (52-90 and 22-76, respectively) experiencing neurodegenerative processes, while the Autism data set features younger patients (2-14). Patterns of normal aging would be expected to differ between these age ranges. Although the COPD controls are in a similar-aged population to the AD and HD controls, we note that the COPD controls are all smokers, the samples are measuring expression in small airway epithelial cells rather than blood, and COPD is not a neurodegenerative disorder. All of these points likely contribute to explain the lack of overlap.

## 4 Conclusions

Many studies have focused on identifying dynamic expression changes in temporal processes. Most of these, however, use either static or traditional time series analyses on self-contained temporal data sets with a limited number of time points (Zinman et al., 2013). Generating additional time points for such analyses involves a cost-benefit tradeoff that has recently been explored (Sefer et al., 2016). Although there is typically greater benefit from adding time points at the expense of replicates, the costs of sampling adequately to identify medically-relevant changes in temporal dynamics may be prohibitive, especially when the dynamic processes are not already well understood.

We have therefore suggested integrating temporal information across static data sets to create a virtual time series, and we introduced an approach based on outlier detection to identify functional pathways or gene sets in which the temporal pattern of expression is disrupted. It is perhaps somewhat surprising that the temporal signal in disease can be strong enough to overcome the noise inherent in combining data points from different subjects, but that observation emphasizes the power to be gained by using an explicit temporal model. Such an approach to data integration will be increasingly valuable as the collections of usable static data in public repositories continue to grow.

This approach may also be applied to any continuous variable, not just time or age, that characterizes high-dimensional data that likely reflects categorical phenotypes or sample characteristics. Potential applications are many, but it seems particularly likely that these methods could be of value in gaining a better mechanistic understanding of developmental disorders or issues in geriatric medicine.

Diseases involving progressive decline or loss of function represent another important application area. Although here we have focused on age as the relevant temporal variable, a more appropriate temporal annotation might be time since diagnosis, or time since some other clinically-defined criterion, rather than age *per se*. It is not then obvious what the appropriate temporal annotation to measure in the control patients would be, but meaningful solutions could be derived for individual use cases. Such an approach might help identify early degenerative or compensatory signals in the course of disease, with potential implications for treatment.

At present, TEMPO identifies only dysregulation in predefined sets of genes. Another important direction for future work is identification of de novo gene sets. Such a method could use the TEMPO dysregulation score as a fitness metric in an optimization algorithm, expanding or recombining known dysregulated gene sets to identify new ones.

Finally, many expression data sets include expression data taken from the same individual at a small number of time points. An interesting and important question for future work is to develop methods for integrating such short time series with static data in an intelligent way. Specifically, the method should make use of dependencies between samples from the same individuals while allowing the use of unrelated samples to learn more about the temporal or age-related expression variation. Doing so will enable better exploitation of existing repositories of transcriptional data for novel discovery.

## Acknowledgements

We thank Diana Bianchi, Jill Maron, and Lystra Hayden for valuable comments on an earlier draft of this manuscript, and members of the Tufts BCB research group for helpful discussions.

## Funding

This work was supported by NIH R01HD076140.

## APPENDIX A

We suspected that, compared to linear regression, PLSR would be better able to handle the dimensions and redundancy of gene expression data (Tobias, 1995). We also considered using Support vector regression (SVR) models with either a linear or radial kernel function (Drucker et al., 1997; Smola and Schölkopf, 2004). On both the ASD autism data set and on an additional developmental data set from GEO (GSE32472), we evaluated the predictive performance of linear regression (LR), PLSR, and SVR models, trained on all control samples using all genes in leave-one-out cross validation. We implemented both LR and SVR in R, the former via the `lm()` function from the `stats` package, the latter in the `e1071` package with default settings.

As hypothesized, linear regression predicted ages less accurately than other methods. On both data sets, PLSR models had lower Mean Squared Error than either SVR model (see Table 4). However, we did not explore the space of possible parameters for SVR.

**Table 4:** Mean squared errors for PLSR, SVR with both linear and radial kernels, and linear models on all control samples in two data sets.

| Method | ASD MSE | GSE32472 MSE |
| --- | --- | --- |
| PLSR | 3.65 | 2.33 |
| SVR (linear kernel) | 4.12 | 4.26 |
| SVR (radial kernel) | 4.29 | 4.25 |
| Linear Regression | 710.74 | 32.72 |

## APPENDIX B

In Section 2.1.2, we model prediction errors using a normal distribution. To test this assumption, we assessed normality using the Shapiro-Wilk normality test in R. On many data sets, we found that the observed error distributions on the control set are in fact normal for nearly all gene sets. However, on some data sets, a substantial fraction of the gene sets have slightly skewed error distributions that do not pass the criteria for normality. We believe that such skewing arises from a lack of uniformity in the age distribution of the control samples.

Some regression models rely on the assumption of normality. However, PLSR is considered relatively robust to data that do not fit this assumption (Farahani et al., 2010). We found that, although there is a modest negative correlation (−0.31) between the normality of the residuals for a gene set and the accuracy of that gene set’s model on the training samples (Figure 6), there are many high-quality models with non-normal residuals.

**Figure 6:**
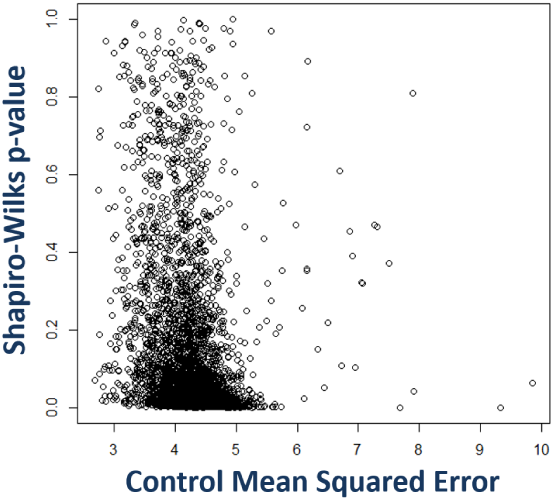
Plot of model quality (control mean squared error) vs. model normality (p-value from the Shapiro-Wilk test, with low p-values *rejecting* normality). Each dot represents a gene set; dots with low MSE and low p-values represent relatively accurate models that fail to meet the criteria for normally distributed residuals.

We emphasize that even in these cases, the non-normality of the prediction errors does not appreciably affect our results. This is because the scoring function does not make use of any specific properties of the normal distribution.

To verify this, we assessed performance of an alternative, nonparametric scoring function:

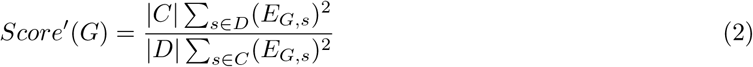

This score is the ratio of the mean squared errors for the disease and control sets. Using this scoring function returns nearly identical ranked lists of gene sets (Spearman rank correlation ≥ .99 between *Score* and *Score′*on all data sets used in this manuscript). We conclude that even if the error distributions are somewhat skewed, the surprisal probabilities are close enough to those expected from the normal distribution that our scoring function is capturing the intended relationship between prediction accuracies in the control and disease sample sets.

## APPENDIX C

**Table 5:**
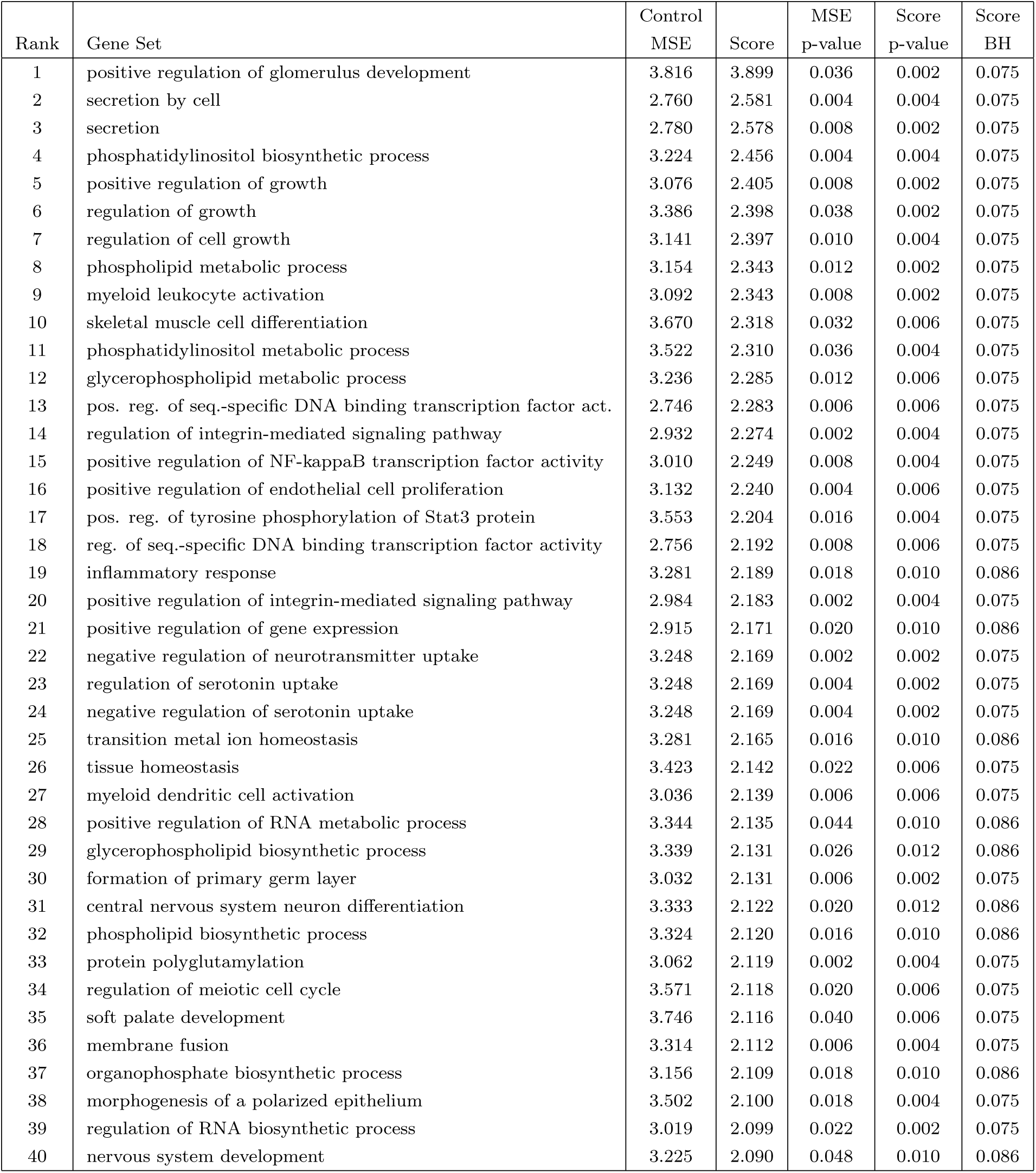
The top 40 highest-scoring gene sets in ASD, ranked by TEMPO score.

| Rank | Gene Set | Control<br>MSE | Score | MSE<br>p-value | Score<br>p-value | Score<br>BH |
| --- | --- | --- | --- | --- | --- | --- |
| 1 | positive regulation of glomerulus development | 3.816 | 3.899 | 0.036 | 0.002 | 0.075 |
| 2 | secretion by cell | 2.760 | 2.581 | 0.004 | 0.004 | 0.075 |
| 3 | secretion | 2.780 | 2.578 | 0.008 | 0.002 | 0.075 |
| 4 | phosphatidylinositol biosynthetic process | 3.224 | 2.456 | 0.004 | 0.004 | 0.075 |
| 5 | positive regulation of growth | 3.076 | 2.405 | 0.008 | 0.002 | 0.075 |
| 6 | regulation of growth | 3.386 | 2.398 | 0.038 | 0.002 | 0.075 |
| 7 | regulation of cell growth | 3.141 | 2.397 | 0.010 | 0.004 | 0.075 |
| 8 | phospholipid metabolic process | 3.154 | 2.343 | 0.012 | 0.002 | 0.075 |
| 9 | myeloid leukocyte activation | 3.092 | 2.343 | 0.008 | 0.002 | 0.075 |
| 10 | skeletal muscle cell differentiation | 3.670 | 2.318 | 0.032 | 0.006 | 0.075 |
| 11 | phosphatidylinositol metabolic process | 3.522 | 2.310 | 0.036 | 0.004 | 0.075 |
| 12 | glycerophospholipid metabolic process | 3.236 | 2.285 | 0.012 | 0.006 | 0.075 |
| 13 | pos. reg. of seq.-specific DNA binding transcription factor act. | 2.746 | 2.283 | 0.006 | 0.006 | 0.075 |
| 14 | regulation of integrin-mediated signaling pathway | 2.932 | 2.274 | 0.002 | 0.004 | 0.075 |
| 15 | positive regulation of NF-kappaB transcription factor activity | 3.010 | 2.249 | 0.008 | 0.004 | 0.075 |
| 16 | positive regulation of endothelial cell proliferation | 3.132 | 2.240 | 0.004 | 0.006 | 0.075 |
| 17 | pos. reg. of tyrosine phosphorylation of Stat3 protein | 3.553 | 2.204 | 0.016 | 0.004 | 0.075 |
| 18 | reg. of seq.-specific DNA binding transcription factor activity | 2.756 | 2.192 | 0.008 | 0.006 | 0.075 |
| 19 | inflammatory response | 3.281 | 2.189 | 0.018 | 0.010 | 0.086 |
| 20 | positive regulation of integrin-mediated signaling pathway | 2.984 | 2.183 | 0.002 | 0.004 | 0.075 |
| 21 | positive regulation of gene expression | 2.915 | 2.171 | 0.020 | 0.010 | 0.086 |
| 22 | negative regulation of neurotransmitter uptake | 3.248 | 2.169 | 0.002 | 0.002 | 0.075 |
| 23 | regulation of serotonin uptake | 3.248 | 2.169 | 0.004 | 0.002 | 0.075 |
| 24 | negative regulation of serotonin uptake | 3.248 | 2.169 | 0.004 | 0.002 | 0.075 |
| 25 | transition metal ion homeostasis | 3.281 | 2.165 | 0.016 | 0.010 | 0.086 |
| 26 | tissue homeostasis | 3.423 | 2.142 | 0.022 | 0.006 | 0.075 |
| 27 | myeloid dendritic cell activation | 3.036 | 2.139 | 0.006 | 0.006 | 0.075 |
| 28 | positive regulation of RNA metabolic process | 3.344 | 2.135 | 0.044 | 0.010 | 0.086 |
| 29 | glycerophospholipid biosynthetic process | 3.339 | 2.131 | 0.026 | 0.012 | 0.086 |
| 30 | formation of primary germ layer | 3.032 | 2.131 | 0.006 | 0.002 | 0.075 |
| 31 | central nervous system neuron differentiation | 3.333 | 2.122 | 0.020 | 0.012 | 0.086 |
| 32 | phospholipid biosynthetic process | 3.324 | 2.120 | 0.016 | 0.010 | 0.086 |
| 33 | protein polyglutamylation | 3.062 | 2.119 | 0.002 | 0.004 | 0.075 |
| 34 | regulation of meiotic cell cycle | 3.571 | 2.118 | 0.020 | 0.006 | 0.075 |
| 35 | soft palate development | 3.746 | 2.116 | 0.040 | 0.006 | 0.075 |
| 36 | membrane fusion | 3.314 | 2.112 | 0.006 | 0.004 | 0.075 |
| 37 | organophosphate biosynthetic process | 3.156 | 2.109 | 0.018 | 0.010 | 0.086 |
| 38 | morphogenesis of a polarized epithelium | 3.502 | 2.100 | 0.018 | 0.004 | 0.075 |
| 39 | regulation of RNA biosynthetic process | 3.019 | 2.099 | 0.022 | 0.002 | 0.075 |
| 40 | nervous system development | 3.225 | 2.090 | 0.048 | 0.010 | 0.086 |

**Table 6:**
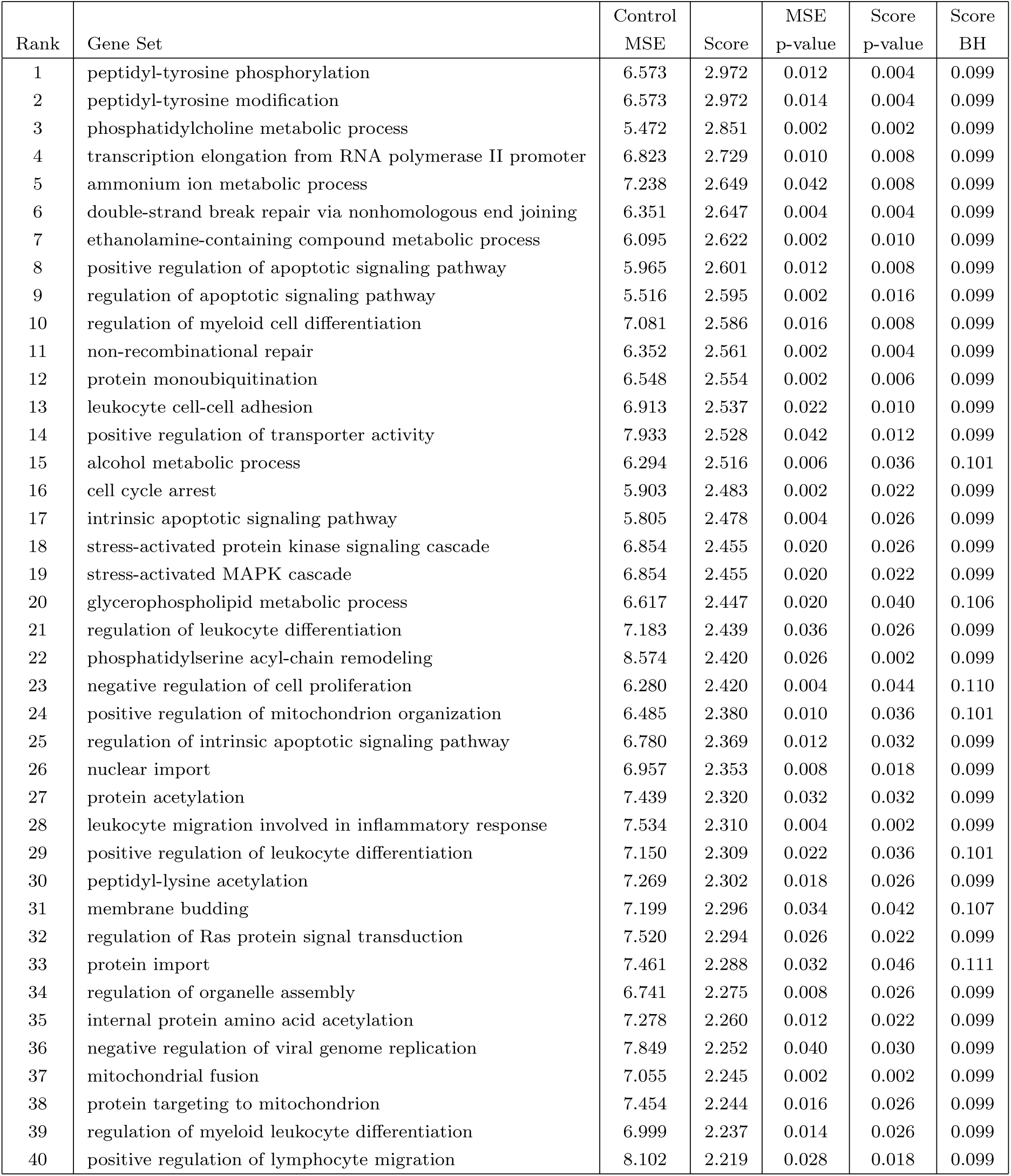
The top 40 highest-scoring gene sets in AD, ranked by TEMPO score.

**Table 7:**
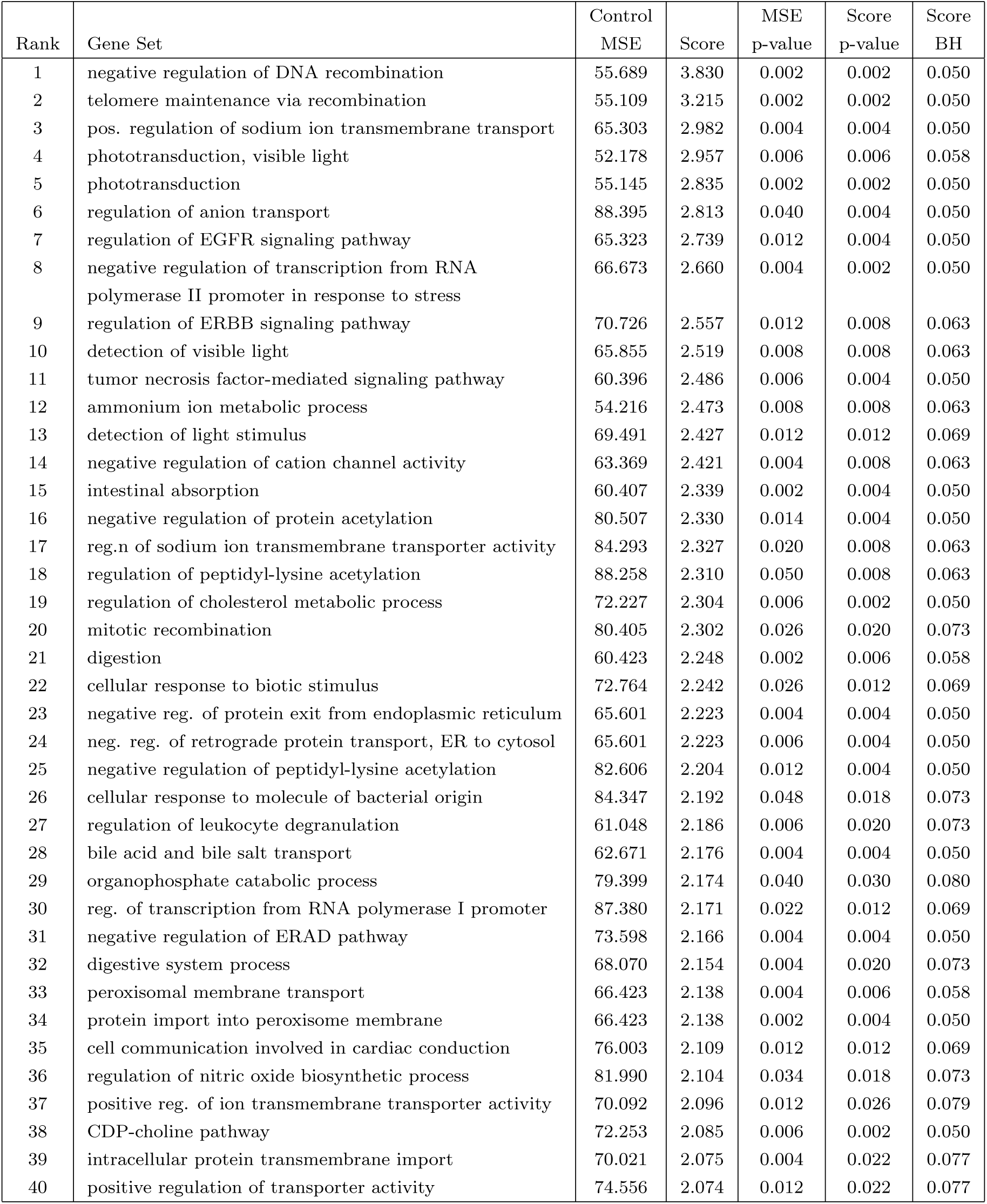
The top 40 highest-scoring gene sets in HD, ranked by TEMPO score.

**Table 8:**
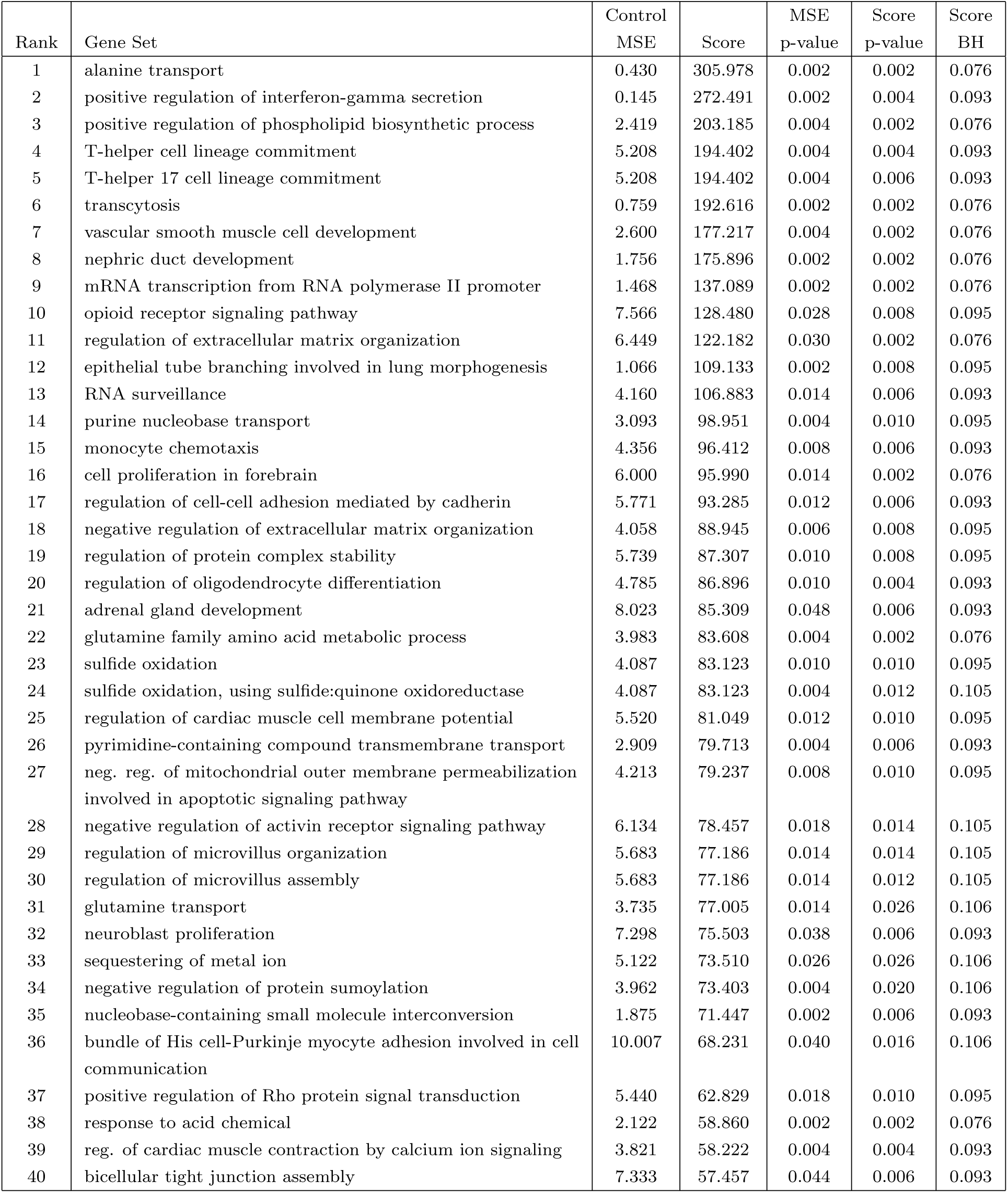
The top 40 highest-scoring gene sets in COPD, ranked by TEMPO score.

## Notes

http://bcb.cs.tufts.edu/tempo

