## Supplemental Tables for "TEMPO: Detecting Pathway-Specific Temporal Dysregulation of Gene Expression in Disease"

### Autism

Table 1: The 40 highest-scoring gene sets ranked by TEMPO score for the ASD data set.

| Rank | Gene Set | Control MSE | Score | MSE p-value | Score p-value | Score BH |
|---|---|---|---|---|---|---|
| 1 | peptidyl-tyrosine phosphorylation | 6.573 | 2.972 | 0.012 | 0.004 | 0.099 |
| 2 | peptidyl-tyrosine modification | 6.573 | 2.972 | 0.014 | 0.004 | 0.099 |
| 3 | phosphatidylcholine metabolic process | 5.472 | 2.851 | 0.002 | 0.002 | 0.099 |
| 4 | transcription elongation from RNA polymerase II promoter | 6.823 | 2.729 | 0.010 | 0.008 | 0.099 |
| 5 | ammonium ion metabolic process | 7.238 | 2.649 | 0.042 | 0.008 | 0.099 |
| 6 | double-strand break repair via nonhomologous end joining | 6.351 | 2.647 | 0.004 | 0.004 | 0.099 |
| 7 | ethanolamine-containing compound metabolic process | 6.095 | 2.622 | 0.002 | 0.010 | 0.099 |
| 8 | positive regulation of apoptotic signaling pathway | 5.965 | 2.601 | 0.012 | 0.008 | 0.099 |
| 9 | regulation of apoptotic signaling pathway | 5.516 | 2.595 | 0.002 | 0.016 | 0.099 |
| 10 | regulation of myeloid cell differentiation | 7.081 | 2.586 | 0.016 | 0.008 | 0.099 |
| 11 | non-recombinational repair | 6.352 | 2.561 | 0.002 | 0.004 | 0.099 |
| 12 | protein monoubiquitination | 6.548 | 2.554 | 0.002 | 0.006 | 0.099 |
| 13 | leukocyte cell-cell adhesion | 6.913 | 2.537 | 0.022 | 0.010 | 0.099 |
| 14 | positive regulation of transporter activity | 7.933 | 2.528 | 0.042 | 0.012 | 0.099 |
| 15 | alcohol metabolic process | 6.294 | 2.516 | 0.006 | 0.036 | 0.101 |
| 16 | cell cycle arrest | 5.903 | 2.483 | 0.002 | 0.022 | 0.099 |
| 17 | intrinsic apoptotic signaling pathway | 5.805 | 2.478 | 0.004 | 0.026 | 0.099 |
| 18 | stress-activated protein kinase signaling cascade | 6.854 | 2.455 | 0.020 | 0.026 | 0.099 |
| 19 | stress-activated MAPK cascade | 6.854 | 2.455 | 0.020 | 0.022 | 0.099 |
| 20 | glycerophospholipid metabolic process | 6.617 | 2.447 | 0.020 | 0.040 | 0.106 |
| 21 | regulation of leukocyte differentiation | 7.183 | 2.439 | 0.036 | 0.026 | 0.099 |
| 22 | phosphatidylserine acyl-chain remodeling | 8.574 | 2.420 | 0.026 | 0.002 | 0.099 |
| 23 | negative regulation of cell proliferation | 6.280 | 2.420 | 0.004 | 0.044 | 0.110 |
| 24 | positive regulation of mitochondrion organization | 6.485 | 2.380 | 0.010 | 0.036 | 0.101 |
| 25 | regulation of intrinsic apoptotic signaling pathway | 6.780 | 2.369 | 0.012 | 0.032 | 0.099 |
| 26 | nuclear import | 6.957 | 2.353 | 0.008 | 0.018 | 0.099 |
| 27 | protein acetylation | 7.439 | 2.320 | 0.032 | 0.032 | 0.099 |
| 28 | leukocyte migration involved in inflammatory response | 7.534 | 2.310 | 0.004 | 0.002 | 0.099 |
| 29 | positive regulation of leukocyte differentiation | 7.150 | 2.309 | 0.022 | 0.036 | 0.101 |
| 30 | peptidyl-lysine acetylation | 7.269 | 2.302 | 0.018 | 0.026 | 0.099 |
| 31 | membrane budding | 7.199 | 2.296 | 0.034 | 0.042 | 0.107 |
| 32 | regulation of Ras protein signal transduction | 7.520 | 2.294 | 0.026 | 0.022 | 0.099 |
| 33 | protein import | 7.461 | 2.288 | 0.032 | 0.046 | 0.111 |
| 34 | regulation of organelle assembly | 6.741 | 2.275 | 0.008 | 0.026 | 0.099 |
| 35 | internal protein amino acid acetylation | 7.278 | 2.260 | 0.012 | 0.022 | 0.099 |
| 36 | negative regulation of viral genome replication | 7.849 | 2.252 | 0.040 | 0.030 | 0.099 |
| 37 | mitochondrial fusion | 7.055 | 2.245 | 0.002 | 0.002 | 0.099 |
| 38 | protein targeting to mitochondrion | 7.454 | 2.244 | 0.016 | 0.026 | 0.099 |
| 39 | regulation of myeloid leukocyte differentiation | 6.999 | 2.237 | 0.014 | 0.026 | 0.099 |
| 40 | positive regulation of lymphocyte migration | 8.102 | 2.219 | 0.028 | 0.018 | 0.099 |

Table 2: The 40 highest-scoring upregulated gene sets returned by GSEA in ASD

| Rank | Gene Set | NES | p-value | FDR |
|---|---|---|---|---|
| 1 | dorsal/ventral axis specification | -1.91 | 0.00 | 1.00 |
| 2 | primary alcohol metabolic process | -1.89 | 0.00 | 1.00 |
| 3 | regulation of chondrocyte differentiation | -1.89 | 0.00 | 0.78 |
| 4 | positive regulation of chondrocyte differentiation | -1.88 | 0.00 | 0.67 |
| 5 | positive regulation of camp biosynthetic process | -1.87 | 0.00 | 0.65 |
| 6 | positive regulation of cyclic nucleotide biosynthetic process | -1.86 | 0.00 | 0.56 |
| 7 | positive regulation of adenylate cyclase activity | -1.86 | 0.00 | 0.49 |
| 8 | organ induction | -1.84 | 0.00 | 0.61 |
| 9 | regulation of carbohydrate biosynthetic process | -1.84 | 0.00 | 0.54 |
| 10 | positive regulation of smoothened signaling pathway | -1.81 | 0.00 | 0.67 |
| 11 | positive regulation of lyase activity | -1.81 | 0.00 | 0.61 |
| 12 | negative regulation of microtubule polymerization | -1.81 | 0.00 | 0.60 |
| 13 | somatic motor neuron differentiation | -1.80 | 0.00 | 0.58 |
| 14 | positive regulation of nucleotide biosynthetic process | -1.80 | 0.00 | 0.57 |
| 15 | positive regulation of purine nucleotide biosynthetic process | -1.80 | 0.00 | 0.53 |
| 16 | modulation by host of viral release from host cell | -1.79 | 0.00 | 0.51 |
| 17 | positive regulation by host of viral release from host cell | -1.79 | 0.00 | 0.48 |
| 18 | phototransduction | -1.79 | 0.00 | 0.47 |
| 19 | negative regulation of t cell apoptotic process | -1.79 | 0.00 | 0.46 |
| 20 | activation of adenylate cyclase activity | -1.79 | 0.00 | 0.44 |
| 21 | negative regulation of platelet-derived growth factor receptor signaling pathway | -1.78 | 0.01 | 0.45 |
| 22 | phototransduction, visible light | -1.77 | 0.00 | 0.47 |
| 23 | notochord development | -1.77 | 0.01 | 0.46 |
| 24 | apical junction assembly | -1.77 | 0.00 | 0.45 |
| 25 | triglyceride catabolic process | -1.76 | 0.01 | 0.47 |
| 26 | developmental induction | -1.76 | 0.00 | 0.48 |
| 27 | regulation of cartilage development | -1.76 | 0.00 | 0.48 |
| 28 | detection of light stimulus | -1.74 | 0.00 | 0.54 |
| 29 | spinal cord development | -1.73 | 0.00 | 0.58 |
| 30 | acid secretion | -1.73 | 0.00 | 0.56 |
| 31 | digestion | -1.73 | 0.00 | 0.55 |
| 32 | positive regulation of protein kinase a signaling | -1.73 | 0.00 | 0.55 |
| 33 | lateral mesoderm development | -1.73 | 0.01 | 0.53 |
| 34 | neutral lipid catabolic process | -1.73 | 0.01 | 0.53 |
| 35 | acylglycerol catabolic process | -1.73 | 0.01 | 0.51 |
| 36 | cyclic nucleotide metabolic process | -1.72 | 0.01 | 0.53 |
| 37 | positive regulation of camp metabolic process | -1.71 | 0.00 | 0.58 |
| 38 | detection of visible light | -1.71 | 0.00 | 0.57 |
| 39 | negative regulation of vascular permeability | -1.71 | 0.01 | 0.57 |
| 40 | aspartate transport | -1.71 | 0.01 | 0.56 |

Table 3: The 40 highest-scoring gene sets returned by maSigPro+GSEA preranked in ASD

| Rank | Gene Set | NES | p-value | FDR |
|---|---|---|---|---|
| 1 | glutaminyl-trnagln biosynthesis via transamidation | 1.48 | 0.00 | 1.00 |
| 2 | vitamin b6 metabolic process | 1.47 | 0.00 | 1.00 |
| 3 | regulation of cortisol secretion | 1.47 | 0.00 | 0.83 |
| 4 | positive regulation of cortisol secretion | 1.47 | 0.00 | 0.64 |
| 5 | positive regulation of glucocorticoid secretion | 1.46 | 0.00 | 0.60 |
| 6 | pyridoxal phosphate metabolic process | 1.46 | 0.01 | 0.54 |
| 7 | cell-cell signaling involved in cell fate commitment | 1.45 | 0.01 | 0.67 |
| 8 | pyridoxal phosphate biosynthetic process | 1.44 | 0.01 | 0.73 |
| 9 | negative regulation of cortisol biosynthetic process | 1.42 | 0.01 | 1.00 |
| 10 | histone h2b conserved c-terminal lysine ubiquitination | 1.42 | 0.01 | 1.00 |
| 11 | negative regulation of tooth mineralization | 1.42 | 0.01 | 1.00 |
| 12 | negative regulation of aldosterone metabolic process | 1.42 | 0.01 | 0.96 |
| 13 | negative regulation of glucocorticoid biosynthetic process | 1.42 | 0.00 | 0.89 |
| 14 | sequestering of triglyceride | 1.42 | 0.00 | 0.84 |
| 15 | negative regulation of aldosterone biosynthetic process | 1.41 | 0.01 | 0.80 |
| 16 | negative regulation of steroid hormone biosynthetic process | 1.41 | 0.00 | 0.81 |
| 17 | aldehyde biosynthetic process | 1.41 | 0.00 | 0.80 |
| 18 | histone h3-k4 demethylation | 1.40 | 0.01 | 0.91 |
| 19 | negative regulation of glucocorticoid metabolic process | 1.40 | 0.01 | 0.87 |
| 20 | oxygen transport | 1.40 | 0.00 | 0.97 |
| 21 | cytoplasmic pattern recognition receptor signaling pathway in response to virus | 1.39 | 0.01 | 0.99 |
| 22 | regulation of toll-like receptor 9 signaling pathway | 1.39 | 0.01 | 1.00 |
| 23 | regulation of glucocorticoid metabolic process | 1.39 | 0.00 | 1.00 |
| 24 | udp-galactose transport | 1.38 | 0.00 | 1.00 |
| 25 | nephric duct morphogenesis | 1.38 | 0.00 | 1.00 |
| 26 | fatty acid beta-oxidation using acyl-coa dehydrogenase | 1.38 | 0.01 | 1.00 |
| 27 | negative regulation of endothelial cell differentiation | 1.37 | 0.02 | 1.00 |
| 28 | regulation of low-density lipoprotein particle clearance | 1.37 | 0.00 | 1.00 |
| 29 | phospholipase c-activating g-protein coupled acetylcholine receptor signaling pathway | 1.37 | 0.00 | 1.00 |
| 30 | regulation of glucocorticoid biosynthetic process | 1.37 | 0.01 | 1.00 |
| 31 | regulation of cortisol biosynthetic process | 1.37 | 0.00 | 1.00 |
| 32 | positive regulation of chaperone-mediated protein complex assembly | 1.37 | 0.02 | 1.00 |
| 33 | negative regulation of epithelial cell differentiation involved in kidney development | 1.37 | 0.00 | 1.00 |
| 34 | negative regulation of histone h3-k36 methylation | 1.36 | 0.01 | 1.00 |
| 35 | negative regulation of nephron tubule epithelial cell differentiation | 1.36 | 0.00 | 1.00 |
| 36 | negative regulation of interleukin-2 secretion | 1.36 | 0.01 | 1.00 |
| 37 | negative regulation of hormone biosynthetic process | 1.36 | 0.00 | 1.00 |
| 38 | regulation of chaperone-mediated protein complex assembly | 1.36 | 0.02 | 1.00 |
| 39 | regulation of type iii interferon production | 1.35 | 0.03 | 1.00 |
| 40 | urinary tract smooth muscle contraction | 1.35 | 0.02 | 1.00 |

### Alzheimer's Disease

Table 4: The 40 highest-scoring gene sets ranked by TEMPO score for the Alzheimer's data set.

| Rank | Gene Set | Control MSE | Score | MSE p-value | Score p-value | Score BH |
|---|---|---|---|---|---|---|
| 1 | peptidyl-tyrosine phosphorylation | 6.573 | 2.972 | 0.012 | 0.004 | 0.099 |
| 2 | peptidyl-tyrosine modification | 6.573 | 2.972 | 0.014 | 0.004 | 0.099 |
| 3 | phosphatidylcholine metabolic process | 5.472 | 2.851 | 0.002 | 0.002 | 0.099 |
| 4 | transcription elongation from RNA polymerase II promoter | 6.823 | 2.729 | 0.010 | 0.008 | 0.099 |
| 5 | ammonium ion metabolic process | 7.238 | 2.649 | 0.042 | 0.008 | 0.099 |
| 6 | double-strand break repair via nonhomologous end joining | 6.351 | 2.647 | 0.004 | 0.004 | 0.099 |
| 7 | ethanolamine-containing compound metabolic process | 6.095 | 2.622 | 0.002 | 0.010 | 0.099 |
| 8 | positive regulation of apoptotic signaling pathway | 5.965 | 2.601 | 0.012 | 0.008 | 0.099 |
| 9 | regulation of apoptotic signaling pathway | 5.516 | 2.595 | 0.002 | 0.016 | 0.099 |
| 10 | regulation of myeloid cell differentiation | 7.081 | 2.586 | 0.016 | 0.008 | 0.099 |
| 11 | non-recombinational repair | 6.352 | 2.561 | 0.002 | 0.004 | 0.099 |
| 12 | protein monoubiquitination | 6.548 | 2.554 | 0.002 | 0.006 | 0.099 |
| 13 | leukocyte cell-cell adhesion | 6.913 | 2.537 | 0.022 | 0.010 | 0.099 |
| 14 | positive regulation of transporter activity | 7.933 | 2.528 | 0.042 | 0.012 | 0.099 |
| 15 | alcohol metabolic process | 6.294 | 2.516 | 0.006 | 0.036 | 0.101 |
| 16 | cell cycle arrest | 5.903 | 2.483 | 0.002 | 0.022 | 0.099 |
| 17 | intrinsic apoptotic signaling pathway | 5.805 | 2.478 | 0.004 | 0.026 | 0.099 |
| 18 | stress-activated protein kinase signaling cascade | 6.854 | 2.455 | 0.020 | 0.026 | 0.099 |
| 19 | stress-activated MAPK cascade | 6.854 | 2.455 | 0.020 | 0.022 | 0.099 |
| 20 | glycerophospholipid metabolic process | 6.617 | 2.447 | 0.020 | 0.040 | 0.106 |
| 21 | regulation of leukocyte differentiation | 7.183 | 2.439 | 0.036 | 0.026 | 0.099 |
| 22 | phosphatidylserine acyl-chain remodeling | 8.574 | 2.420 | 0.026 | 0.002 | 0.099 |
| 23 | negative regulation of cell proliferation | 6.280 | 2.420 | 0.004 | 0.044 | 0.110 |
| 24 | positive regulation of mitochondrion organization | 6.485 | 2.380 | 0.010 | 0.036 | 0.101 |
| 25 | regulation of intrinsic apoptotic signaling pathway | 6.780 | 2.369 | 0.012 | 0.032 | 0.099 |
| 26 | nuclear import | 6.957 | 2.353 | 0.008 | 0.018 | 0.099 |
| 27 | protein acetylation | 7.439 | 2.320 | 0.032 | 0.032 | 0.099 |
| 28 | leukocyte migration involved in inflammatory response | 7.534 | 2.310 | 0.004 | 0.002 | 0.099 |
| 29 | positive regulation of leukocyte differentiation | 7.150 | 2.309 | 0.022 | 0.036 | 0.101 |
| 30 | peptidyl-lysine acetylation | 7.269 | 2.302 | 0.018 | 0.026 | 0.099 |
| 31 | membrane budding | 7.199 | 2.296 | 0.034 | 0.042 | 0.107 |
| 32 | regulation of Ras protein signal transduction | 7.520 | 2.294 | 0.026 | 0.022 | 0.099 |
| 33 | protein import | 7.461 | 2.288 | 0.032 | 0.046 | 0.111 |
| 34 | regulation of organelle assembly | 6.741 | 2.275 | 0.008 | 0.026 | 0.099 |
| 35 | internal protein amino acid acetylation | 7.278 | 2.260 | 0.012 | 0.022 | 0.099 |
| 36 | negative regulation of viral genome replication | 7.849 | 2.252 | 0.040 | 0.030 | 0.099 |
| 37 | mitochondrial fusion | 7.055 | 2.245 | 0.002 | 0.002 | 0.099 |
| 38 | protein targeting to mitochondrion | 7.454 | 2.244 | 0.016 | 0.026 | 0.099 |
| 39 | regulation of myeloid leukocyte differentiation | 6.999 | 2.237 | 0.014 | 0.026 | 0.099 |
| 40 | positive regulation of lymphocyte migration | 8.102 | 2.219 | 0.028 | 0.018 | 0.099 |

Table 5: The 40 highest-scoring gene sets returned by GSEA in Alzheimer's

| Rank | Gene Set | NES | p-value | FDR |
|---|---|---|---|---|
| 1 | transepithelial transport | -2.064 | 0.000 | 0.025 |
| 2 | neuron cell-cell adhesion | -1.929 | 0.000 | 0.135 |
| 3 | inorganic anion transport | -1.921 | 0.000 | 0.106 |
| 4 | cardiac ventricle morphogenesis | -1.916 | 0.000 | 0.087 |
| 5 | neuron migration | -1.915 | 0.000 | 0.071 |
| 6 | regulation of smad protein import into nucleus | -1.893 | 0.002 | 0.078 |
| 7 | positive regulation of glycogen metabolic process | -1.885 | 0.000 | 0.076 |
| 8 | chondroitin sulfate proteoglycan metabolic process | -1.881 | 0.000 | 0.071 |
| 9 | cardiac muscle tissue morphogenesis | -1.869 | 0.000 | 0.077 |
| 10 | mesenchyme morphogenesis | -1.864 | 0.000 | 0.075 |
| 11 | chloride transport | -1.861 | 0.000 | 0.071 |
| 12 | chondroitin sulfate metabolic process | -1.853 | 0.000 | 0.075 |
| 13 | cyclic nucleotide metabolic process | -1.851 | 0.002 | 0.071 |
| 14 | endocardial cushion formation | -1.851 | 0.000 | 0.066 |
| 15 | mucopolysaccharide metabolic process | -1.847 | 0.000 | 0.065 |
| 16 | positive regulation of glucose metabolic process | -1.843 | 0.000 | 0.065 |
| 17 | muscle organ morphogenesis | -1.837 | 0.000 | 0.066 |
| 18 | muscle tissue morphogenesis | -1.837 | 0.000 | 0.062 |
| 19 | regulation of cholesterol storage | -1.833 | 0.000 | 0.062 |
| 20 | regulation of glycogen metabolic process | -1.833 | 0.000 | 0.059 |
| 21 | sodium-independent organic anion transport | -1.829 | 0.000 | 0.059 |
| 22 | glycosaminoglycan metabolic process | -1.827 | 0.000 | 0.059 |
| 23 | regulation of synapse assembly | -1.822 | 0.004 | 0.060 |
| 24 | aminoglycan metabolic process | -1.817 | 0.000 | 0.063 |
| 25 | positive regulation of glycogen biosynthetic process | -1.814 | 0.000 | 0.061 |
| 26 | pulmonary valve development | -1.810 | 0.000 | 0.063 |
| 27 | pulmonary valve morphogenesis | -1.810 | 0.000 | 0.061 |
| 28 | chloride transmembrane transport | -1.808 | 0.000 | 0.060 |
| 29 | cardioblast differentiation | -1.807 | 0.002 | 0.058 |
| 30 | negative regulation of peptide hormone secretion | -1.799 | 0.000 | 0.061 |
| 31 | inorganic anion transmembrane transport | -1.799 | 0.000 | 0.060 |
| 32 | positive regulation of cardiac muscle tissue development | -1.797 | 0.000 | 0.059 |
| 33 | glycosaminoglycan biosynthetic process | -1.795 | 0.000 | 0.059 |
| 34 | positive regulation of lipid transport | -1.793 | 0.000 | 0.060 |
| 35 | ventricular cardiac muscle tissue development | -1.785 | 0.000 | 0.065 |
| 36 | blood vessel endothelial cell differentiation | -1.785 | 0.000 | 0.063 |
| 37 | camp metabolic process | -1.784 | 0.006 | 0.062 |
| 38 | excretion | -1.784 | 0.002 | 0.060 |
| 39 | polyol transport | -1.783 | 0.000 | 0.060 |
| 40 | aminoglycan biosynthetic process | -1.781 | 0.000 | 0.060 |

Table 6: The 40 highest-scoring gene sets returned by maSigPro+GSEA preranked in Alzheimer's

| Rank | Gene Set | NES | p-value | FDR |
|---|---|---|---|---|
| 1 | equilibrioception | 1.510 | 0.000 | 0.686 |
| 2 | aortic valve morphogenesis | 1.478 | 0.000 | 1.000 |
| 3 | aortic valve development | 1.469 | 0.001 | 0.976 |
| 4 | alkaloid metabolic process | 1.429 | 0.012 | 1.000 |
| 5 | intracellular transport of viral protein in host cell | 1.427 | 0.006 | 1.000 |
| 6 | intracellular protein transport in other organism involved in symbiotic interaction | 1.424 | 0.008 | 1.000 |
| 7 | heterochromatin organization | 1.421 | 0.001 | 1.000 |
| 8 | symbiont intracellular protein transport in host | 1.420 | 0.002 | 1.000 |
| 9 | positive regulation of organic acid transport | 1.419 | 0.005 | 1.000 |
| 10 | atp synthesis coupled proton transport | 1.418 | 0.000 | 1.000 |
| 11 | energy coupled proton transport, down electrochemical gradient | 1.417 | 0.000 | 1.000 |
| 12 | mitochondrial atp synthesis coupled proton transport | 1.415 | 0.000 | 0.986 |
| 13 | wound healing, spreading of epidermal cells | 1.409 | 0.008 | 1.000 |
| 14 | regulation of axon extension involved in axon guidance | 1.408 | 0.006 | 0.993 |
| 15 | electron transport chain | 1.407 | 0.000 | 0.951 |
| 16 | oxygen homeostasis | 1.406 | 0.007 | 0.911 |
| 17 | positive regulation of icosanoid secretion | 1.404 | 0.011 | 0.877 |
| 18 | positive regulation of axon guidance | 1.402 | 0.008 | 0.877 |
| 19 | respiratory electron transport chain | 1.402 | 0.000 | 0.833 |
| 20 | camp-mediated signaling | 1.401 | 0.000 | 0.793 |
| 21 | spermatid nucleus differentiation | 1.400 | 0.005 | 0.773 |
| 22 | positive regulation of t cell differentiation in thymus | 1.399 | 0.005 | 0.752 |
| 23 | sperm chromatin condensation | 1.398 | 0.012 | 0.732 |
| 24 | negative regulation of oxidative stress-induced neuron intrinsic apoptotic signaling pathway | 1.398 | 0.009 | 0.704 |
| 25 | myelin assembly | 1.397 | 0.009 | 0.694 |
| 26 | positive regulation of transcription from rna polymerase ii promoter involved in heart development | 1.395 | 0.015 | 0.696 |
| 27 | cardiac chamber formation | 1.395 | 0.010 | 0.672 |
| 28 | heterochromatin assembly | 1.394 | 0.005 | 0.658 |
| 29 | positive regulation of axon extension involved in axon guidance | 1.394 | 0.007 | 0.639 |
| 30 | negative regulation of jun kinase activity | 1.392 | 0.006 | 0.634 |
| 31 | positive regulation of fatty acid transport | 1.391 | 0.006 | 0.627 |
| 32 | positive regulation of fibroblast apoptotic process | 1.389 | 0.011 | 0.632 |
| 33 | positive regulation of thymocyte aggregation | 1.384 | 0.006 | 0.673 |
| 34 | nls-bearing protein import into nucleus | 1.383 | 0.000 | 0.664 |
| 35 | camera-type eye photoreceptor cell differentiation | 1.379 | 0.013 | 0.702 |
| 36 | negative regulation of hydrogen peroxide-mediated programmed cell death | 1.379 | 0.017 | 0.684 |
| 37 | tissue regeneration | 1.377 | 0.007 | 0.685 |
| 38 | riboflavin metabolic process | 1.366 | 0.015 | 0.806 |
| 39 | translational elongation | 1.366 | 0.000 | 0.786 |
| 40 | regulation of rna export from nucleus | 1.366 | 0.006 | 0.774 |

### Huntington's Disease

Table 7: The 40 highest-scoring gene sets ranked by TEMPO score for the Huntington's Disease data set.

| Rank | Gene Set | Control MSE | Score | MSE p-value | Score p-value | Score BH |
|---|---|---|---|---|---|---|
| 1 | negative regulation of DNA recombination | 55.689 | 3.830 | 0.002 | 0.002 | 0.050 |
| 2 | telomere maintenance via recombination | 55.109 | 3.215 | 0.002 | 0.002 | 0.050 |
| 3 | pos. regulation of sodium ion transmembrane transport | 65.303 | 2.982 | 0.004 | 0.004 | 0.050 |
| 4 | phototransduction, visible light | 52.178 | 2.957 | 0.006 | 0.006 | 0.058 |
| 5 | phototransduction | 55.145 | 2.835 | 0.002 | 0.002 | 0.050 |
| 6 | regulation of anion transport | 88.395 | 2.813 | 0.040 | 0.004 | 0.050 |
| 7 | regulation of EGFR signaling pathway | 65.323 | 2.739 | 0.012 | 0.004 | 0.050 |
| 8 | negative regulation of transcription from RNA polymerase II promoter in response to stress | 66.673 | 2.660 | 0.004 | 0.002 | 0.050 |
| 9 | regulation of ERBB signaling pathway | 70.726 | 2.557 | 0.012 | 0.008 | 0.063 |
| 10 | detection of visible light | 65.855 | 2.519 | 0.008 | 0.008 | 0.063 |
| 11 | tumor necrosis factor-mediated signaling pathway | 60.396 | 2.486 | 0.006 | 0.004 | 0.050 |
| 12 | ammonium ion metabolic process | 54.216 | 2.473 | 0.008 | 0.008 | 0.063 |
| 13 | detection of light stimulus | 69.491 | 2.427 | 0.012 | 0.012 | 0.069 |
| 14 | negative regulation of cation channel activity | 63.369 | 2.421 | 0.004 | 0.008 | 0.063 |
| 15 | intestinal absorption | 60.407 | 2.339 | 0.002 | 0.004 | 0.050 |
| 16 | negative regulation of protein acetylation | 80.507 | 2.330 | 0.014 | 0.004 | 0.050 |
| 17 | reg.n of sodium ion transmembrane transporter activity | 84.293 | 2.327 | 0.020 | 0.008 | 0.063 |
| 18 | regulation of peptidyl-lysine acetylation | 88.258 | 2.310 | 0.050 | 0.008 | 0.063 |
| 19 | regulation of cholesterol metabolic process | 72.227 | 2.304 | 0.006 | 0.002 | 0.050 |
| 20 | mitotic recombination | 80.405 | 2.302 | 0.026 | 0.020 | 0.073 |
| 21 | digestion | 60.423 | 2.248 | 0.002 | 0.006 | 0.058 |
| 22 | cellular response to biotic stimulus | 72.764 | 2.242 | 0.026 | 0.012 | 0.069 |
| 23 | negative reg. of protein exit from endoplasmic reticulum | 65.601 | 2.223 | 0.004 | 0.004 | 0.050 |
| 24 | neg. reg. of retrograde protein transport, ER to cytosol | 65.601 | 2.223 | 0.006 | 0.004 | 0.050 |
| 25 | negative regulation of peptidyl-lysine acetylation | 82.606 | 2.204 | 0.012 | 0.004 | 0.050 |
| 26 | cellular response to molecule of bacterial origin | 84.347 | 2.192 | 0.048 | 0.018 | 0.073 |
| 27 | regulation of leukocyte degranulation | 61.048 | 2.186 | 0.006 | 0.020 | 0.073 |
| 28 | bile acid and bile salt transport | 62.671 | 2.176 | 0.004 | 0.004 | 0.050 |
| 29 | organophosphate catabolic process | 79.399 | 2.174 | 0.040 | 0.030 | 0.080 |
| 30 | reg. of transcription from RNA polymerase I promoter | 87.380 | 2.171 | 0.022 | 0.012 | 0.069 |
| 31 | negative regulation of ERAD pathway | 73.598 | 2.166 | 0.004 | 0.004 | 0.050 |
| 32 | digestive system process | 68.070 | 2.154 | 0.004 | 0.020 | 0.073 |
| 33 | peroxisomal membrane transport | 66.423 | 2.138 | 0.004 | 0.006 | 0.058 |
| 34 | protein import into peroxisome membrane | 66.423 | 2.138 | 0.002 | 0.004 | 0.050 |
| 35 | cell communication involved in cardiac conduction | 76.003 | 2.109 | 0.012 | 0.012 | 0.069 |
| 36 | regulation of nitric oxide biosynthetic process | 81.990 | 2.104 | 0.034 | 0.018 | 0.073 |
| 37 | positive reg. of ion transmembrane transporter activity | 70.092 | 2.096 | 0.012 | 0.026 | 0.079 |
| 38 | CDP-choline pathway | 72.253 | 2.085 | 0.006 | 0.002 | 0.050 |
| 39 | intracellular protein transmembrane import | 70.021 | 2.075 | 0.004 | 0.022 | 0.077 |
| 40 | positive regulation of transporter activity | 74.556 | 2.074 | 0.012 | 0.022 | 0.077 |

Table 8: The 40 highest-scoring gene sets ranked by TEMPO score for the Huntington's Disease data set, with symptomatic patients only.

| Rank | Gene Set | Control MSE | Score | MSE p-value | Score p-value | Score BH |
|---|---|---|---|---|---|---|
| 1 | negative regulation of DNA recombination | 55.689 | 4.273 | 0.002 | 0.002 | 0.043 |
| 2 | positive regulation of sodium ion transmembrane transport | 65.303 | 3.452 | 0.002 | 0.002 | 0.043 |
| 3 | telomere maintenance via recombination | 55.109 | 3.422 | 0.002 | 0.004 | 0.047 |
| 4 | phototransduction, visible light | 52.178 | 3.375 | 0.002 | 0.002 | 0.043 |
| 5 | phototransduction | 55.145 | 3.215 | 0.002 | 0.002 | 0.043 |
| 6 | negative regulation of transcription from RNA polymerase II promoter in response to stress | 66.673 | 3.070 | 0.004 | 0.002 | 0.043 |
| 7 | regulation of EGFR signaling pathway | 65.323 | 3.039 | 0.012 | 0.002 | 0.043 |
| 8 | tumor necrosis factor-mediated signaling pathway | 60.396 | 2.894 | 0.012 | 0.002 | 0.043 |
| 9 | detection of visible light | 65.855 | 2.886 | 0.014 | 0.006 | 0.059 |
| 10 | regulation of ERBB signaling pathway | 70.726 | 2.843 | 0.008 | 0.004 | 0.047 |
| 11 | ammonium ion metabolic process | 54.216 | 2.782 | 0.002 | 0.004 | 0.047 |
| 12 | detection of light stimulus | 69.491 | 2.767 | 0.014 | 0.012 | 0.073 |
| 13 | regulation of sodium ion transmembrane transporter activity | 84.293 | 2.719 | 0.030 | 0.004 | 0.047 |
| 14 | intestinal absorption | 60.407 | 2.665 | 0.002 | 0.004 | 0.047 |
| 15 | negative regulation of cation channel activity | 63.369 | 2.638 | 0.002 | 0.002 | 0.043 |
| 16 | digestion | 60.423 | 2.595 | 0.004 | 0.012 | 0.073 |
| 17 | negative regulation of protein acetylation | 80.507 | 2.552 | 0.016 | 0.006 | 0.059 |
| 18 | regulation of peptidyl-lysine acetylation | 88.258 | 2.538 | 0.048 | 0.010 | 0.070 |
| 19 | bile acid and bile salt transport | 62.671 | 2.528 | 0.002 | 0.006 | 0.059 |
| 20 | regulation of cholesterol metabolic process | 72.227 | 2.517 | 0.004 | 0.002 | 0.043 |
| 21 | digestive system process | 68.070 | 2.504 | 0.006 | 0.008 | 0.064 |
| 22 | reg. of transcription from RNAP II promoter in response to stress | 86.002 | 2.465 | 0.036 | 0.008 | 0.064 |
| 23 | cellular response to biotic stimulus | 72.764 | 2.453 | 0.008 | 0.008 | 0.064 |
| 24 | mitotic recombination | 80.405 | 2.447 | 0.016 | 0.022 | 0.077 |
| 25 | negative regulation of peptidyl-lysine acetylation | 82.606 | 2.437 | 0.012 | 0.008 | 0.064 |
| 26 | positive regulation of transporter activity | 74.556 | 2.417 | 0.014 | 0.014 | 0.074 |
| 27 | organophosphate catabolic process | 79.399 | 2.408 | 0.038 | 0.022 | 0.077 |
| 28 | positive regulation of ion transmembrane transporter activity | 70.092 | 2.405 | 0.006 | 0.018 | 0.077 |
| 29 | regulation of transcription from RNA polymerase I promoter | 87.380 | 2.403 | 0.026 | 0.014 | 0.074 |
| 30 | locomotor rhythm | 64.605 | 2.382 | 0.002 | 0.002 | 0.043 |
| 31 | peroxisomal membrane transport | 66.423 | 2.381 | 0.004 | 0.004 | 0.047 |
| 32 | protein import into peroxisome membrane | 66.423 | 2.381 | 0.002 | 0.002 | 0.043 |
| 33 | neg. regulation of protein exit from endoplasmic reticulum | 65.601 | 2.356 | 0.006 | 0.016 | 0.077 |
| 34 | neg. regulation of retrograde protein transport, ER to cytosol | 65.601 | 2.356 | 0.002 | 0.002 | 0.043 |
| 35 | positive regulation of cation transmembrane transport | 68.454 | 2.348 | 0.010 | 0.018 | 0.077 |
| 36 | synapse organization | 91.797 | 2.342 | 0.048 | 0.018 | 0.077 |
| 37 | cell communication involved in cardiac conduction | 76.003 | 2.337 | 0.012 | 0.014 | 0.074 |
| 38 | blood coagulation, intrinsic pathway | 77.824 | 2.324 | 0.014 | 0.004 | 0.047 |
| 39 | cellular aldehyde metabolic process | 79.671 | 2.285 | 0.024 | 0.020 | 0.077 |
| 40 | negative regulation of ERAD pathway | 73.598 | 2.279 | 0.010 | 0.012 | 0.073 |

Table 9: The 40 highest-scoring gene sets ranked by TEMPO score for the Huntington's Disease data set, with pre-symptomatic patients only.

| Rank | Gene Set | Control MSE | Score | MSE p-value | Score p-value | Score BH |
|---|---|---|---|---|---|---|
| 1 | negative regulation of DNA recombination | 55.689 | 2.778 | 0.002 | 0.002 | 0.051 |
| 2 | telomere maintenance via recombination | 55.109 | 2.725 | 0.002 | 0.002 | 0.051 |
| 3 | regulation of anion transport | 88.395 | 2.418 | 0.042 | 0.006 | 0.068 |
| 4 | neg. reg. of reactive oxygen species biosynthetic process | 84.943 | 2.104 | 0.004 | 0.002 | 0.051 |
| 5 | regulation of nitric oxide biosynthetic process | 81.990 | 2.091 | 0.018 | 0.006 | 0.068 |
| 6 | regulation of EGFR signaling pathway | 65.323 | 2.027 | 0.010 | 0.004 | 0.068 |
| 7 | intracellular protein transmembrane import | 70.021 | 2.000 | 0.004 | 0.010 | 0.075 |
| 8 | regulation of leukocyte degranulation | 61.048 | 1.970 | 0.004 | 0.012 | 0.075 |
| 9 | phototransduction, visible light | 52.178 | 1.967 | 0.002 | 0.012 | 0.075 |
| 10 | mitotic recombination | 80.405 | 1.960 | 0.024 | 0.010 | 0.075 |
| 11 | intracellular protein transmembrane transport | 78.604 | 1.937 | 0.014 | 0.008 | 0.070 |
| 12 | phototransduction | 55.145 | 1.934 | 0.002 | 0.006 | 0.068 |
| 13 | negative regulation of telomere capping | 85.164 | 1.915 | 0.008 | 0.002 | 0.051 |
| 14 | negative regulation of cation channel activity | 63.369 | 1.906 | 0.010 | 0.010 | 0.075 |
| 15 | neg. regulation of protein exit from endoplasmic reticulum | 65.601 | 1.906 | 0.002 | 0.002 | 0.051 |
| 16 | neg. regulation of retrograde protein transport, ER to cytosol | 65.601 | 1.906 | 0.002 | 0.002 | 0.051 |
| 17 | negative regulation of ERAD pathway | 73.598 | 1.898 | 0.010 | 0.006 | 0.068 |
| 18 | regulation of ERBB signaling pathway | 70.726 | 1.880 | 0.020 | 0.014 | 0.075 |
| 19 | positive regulation of sodium ion transmembrane transport | 65.303 | 1.869 | 0.002 | 0.004 | 0.068 |
| 20 | cellular glucose homeostasis | 83.696 | 1.852 | 0.020 | 0.008 | 0.070 |
| 21 | negative regulation of protein acetylation | 80.507 | 1.803 | 0.012 | 0.006 | 0.068 |
| 22 | regulation of cholesterol metabolic process | 72.227 | 1.799 | 0.008 | 0.006 | 0.068 |
| 23 | mitotic G2/M transition checkpoint | 86.450 | 1.795 | 0.024 | 0.016 | 0.077 |
| 24 | cellular response to biotic stimulus | 72.764 | 1.741 | 0.022 | 0.020 | 0.079 |
| 25 | ammonium ion metabolic process | 54.216 | 1.739 | 0.004 | 0.014 | 0.075 |
| 26 | DNA strand elongation | 89.665 | 1.703 | 0.048 | 0.026 | 0.082 |
| 27 | negative regulation of histone H3-K9 methylation | 86.229 | 1.695 | 0.014 | 0.002 | 0.051 |
| 28 | negative regulation of transcription from RNA polymerase II promoter in response to stress | 66.673 | 1.687 | 0.002 | 0.008 | 0.070 |
| 29 | phosphatidylethanolamine acyl-chain remodeling | 84.213 | 1.665 | 0.016 | 0.008 | 0.070 |
| 30 | protein heterooligomerization | 92.321 | 1.662 | 0.034 | 0.018 | 0.077 |
| 31 | negative regulation of peptidyl-lysine acetylation | 82.606 | 1.650 | 0.012 | 0.020 | 0.079 |
| 32 | detection of visible light | 65.855 | 1.649 | 0.016 | 0.028 | 0.082 |
| 33 | pos. regulation of potassium ion transmembrane transport | 80.479 | 1.637 | 0.016 | 0.014 | 0.075 |
| 34 | developmental process involved in reproduction | 81.601 | 1.633 | 0.026 | 0.008 | 0.070 |
| 35 | acute inflammatory response | 75.302 | 1.633 | 0.010 | 0.024 | 0.082 |
| 36 | CDP-choline pathway | 72.253 | 1.632 | 0.002 | 0.002 | 0.051 |
| 37 | negative regulation of nitric oxide biosynthetic process | 96.945 | 1.629 | 0.048 | 0.002 | 0.051 |
| 38 | negative regulation of nitric oxide metabolic process | 96.945 | 1.629 | 0.044 | 0.002 | 0.051 |
| 39 | protein targeting to peroxisome | 68.215 | 1.629 | 0.012 | 0.024 | 0.082 |
| 40 | peroxisomal transport | 68.215 | 1.629 | 0.008 | 0.026 | 0.082 |

Table 10: The 40 highest-scoring gene sets returned by GSEA in Huntingtons

| Rank | Gene Set | NES | p-value | FDR |
|---|---|---|---|---|
| 1 | regulation of symbiosis, encompassing mutualism through parasitism | -1.87 | 0.00 | 1.00 |
| 2 | carnitine shuttle | -1.84 | 0.00 | 1.00 |
| 3 | negative regulation of insulin receptor signaling pathway | -1.83 | 0.01 | 1.00 |
| 4 | regulation of insulin receptor signaling pathway | -1.81 | 0.00 | 1.00 |
| 5 | negative regulation of cellular response to insulin stimulus | -1.76 | 0.01 | 1.00 |
| 6 | amino-acid betaine transport | -1.75 | 0.01 | 1.00 |
| 7 | carnitine transport | -1.75 | 0.01 | 1.00 |
| 8 | bone morphogenesis | -1.73 | 0.01 | 1.00 |
| 9 | sarcomere organization | -1.73 | 0.01 | 1.00 |
| 10 | endochondral ossification | -1.73 | 0.00 | 1.00 |
| 11 | replacement ossification | -1.73 | 0.00 | 1.00 |
| 12 | regulation of wound healing | -1.71 | 0.00 | 1.00 |
| 13 | positive regulation of leukocyte chemotaxis | -1.71 | 0.02 | 1.00 |
| 14 | actin-mediated cell contraction | -1.70 | 0.01 | 1.00 |
| 15 | fatty acid transmembrane transport | -1.69 | 0.01 | 1.00 |
| 16 | positive regulation of cell-substrate adhesion | -1.68 | 0.04 | 1.00 |
| 17 | anterior/posterior axis specification | -1.68 | 0.01 | 1.00 |
| 18 | regulation of blood pressure | -1.67 | 0.00 | 1.00 |
| 19 | positive regulation of cell-matrix adhesion | -1.66 | 0.03 | 1.00 |
| 20 | activation of mapkk activity | -1.65 | 0.02 | 1.00 |
| 21 | positive regulation of endothelial cell proliferation | -1.65 | 0.01 | 1.00 |
| 22 | phosphatidylglycerol biosynthetic process | -1.65 | 0.03 | 1.00 |
| 23 | nucleoside bisphosphate metabolic process | -1.64 | 0.01 | 1.00 |
| 24 | ribonucleoside bisphosphate metabolic process | -1.64 | 0.01 | 1.00 |
| 25 | purine nucleoside bisphosphate metabolic process | -1.64 | 0.01 | 1.00 |
| 26 | regulation of establishment or maintenance of cell polarity | -1.64 | 0.02 | 1.00 |
| 27 | purine ribonucleoside bisphosphate metabolic process | -1.64 | 0.01 | 1.00 |
| 28 | 3'-phosphoadenosine 5'-phosphosulfate metabolic process | -1.64 | 0.01 | 1.00 |
| 29 | copper ion transport | -1.64 | 0.03 | 1.00 |
| 30 | bone development | -1.64 | 0.01 | 1.00 |
| 31 | regulation of coagulation | -1.63 | 0.01 | 1.00 |
| 32 | regulation of blood coagulation | -1.63 | 0.02 | 1.00 |
| 33 | regulation of hemostasis | -1.63 | 0.02 | 1.00 |
| 34 | n-acetylneuraminate metabolic process | -1.62 | 0.01 | 1.00 |
| 35 | inflammatory response | -1.62 | 0.02 | 1.00 |
| 36 | neutrophil activation | -1.62 | 0.04 | 1.00 |
| 37 | endochondral bone morphogenesis | -1.61 | 0.02 | 1.00 |
| 38 | regulated secretory pathway | -1.61 | 0.03 | 1.00 |
| 39 | post-embryonic hemopoiesis | -1.61 | 0.00 | 1.00 |
| 40 | trabecula formation | -1.61 | 0.02 | 1.00 |

Table 11: The 40 highest-scoring gene sets returned by maSigPro+GSEA preranked in Huntingtons

| Rank | Gene Set | NES | p-value | FDR |
|---|---|---|---|---|
| 1 | steroid biosynthetic process | 2.31 | 0.00 | 0.71 |
| 2 | dna metabolic process | 2.12 | 0.00 | 1.00 |
| 3 | transcription-coupled nucleotide-excision repair | 2.11 | 0.01 | 1.00 |
| 4 | negative regulation of rna metabolic process | 2.11 | 0.00 | 0.96 |
| 5 | cellular response to reactive oxygen species | 2.07 | 0.00 | 0.97 |
| 6 | dna strand elongation involved in dna replication | 2.05 | 0.00 | 1.00 |
| 7 | negative regulation of nitrogen compound metabolic process | 2.04 | 0.00 | 0.87 |
| 8 | response to oxidative stress | 2.04 | 0.00 | 0.77 |
| 9 | cholesterol biosynthetic process | 2.03 | 0.00 | 0.73 |
| 10 | peroxisome fission | 2.02 | 0.00 | 0.72 |
| 11 | deoxyribose phosphate metabolic process | 2.01 | 0.00 | 0.69 |
| 12 | dna repair | 2.01 | 0.00 | 0.65 |
| 13 | cgmp metabolic process | 2.01 | 0.00 | 0.61 |
| 14 | sterol biosynthetic process | 2.00 | 0.00 | 0.58 |
| 15 | positive regulation of gtpase activity | 2.00 | 0.01 | 0.54 |
| 16 | regulation of dna-dependent transcription in response to stress | 2.00 | 0.00 | 0.52 |
| 17 | monocyte differentiation | 1.98 | 0.00 | 0.56 |
| 18 | regulation of histone h3-k4 methylation | 1.98 | 0.00 | 0.53 |
| 19 | negative regulation of transcription, dna-dependent | 1.98 | 0.00 | 0.51 |
| 20 | regulation of cytokine production involved in immune response | 1.97 | 0.00 | 0.50 |
| 21 | metabolic process | 1.97 | 0.01 | 0.48 |
| 22 | alcohol biosynthetic process | 1.97 | 0.01 | 0.47 |
| 23 | deoxyribonucleotide catabolic process | 1.96 | 0.01 | 0.46 |
| 24 | cholesterol metabolic process | 1.96 | 0.01 | 0.45 |
| 25 | regulation of nucleoside metabolic process | 1.95 | 0.01 | 0.46 |
| 26 | nucleotide-excision repair | 1.94 | 0.01 | 0.47 |
| 27 | negative regulation of immune effector process | 1.94 | 0.01 | 0.45 |
| 28 | regulation of transcription from rna polymerase ii promoter in response to stress | 1.94 | 0.01 | 0.44 |
| 29 | negative regulation of nucleobase-containing compound metabolic process | 1.94 | 0.00 | 0.43 |
| 30 | response to reactive oxygen species | 1.93 | 0.01 | 0.43 |
| 31 | deoxyribose phosphate catabolic process | 1.93 | 0.00 | 0.42 |
| 32 | protein k48-linked ubiquitination | 1.93 | 0.01 | 0.41 |
| 33 | regulation of ras protein signal transduction | 1.93 | 0.01 | 0.40 |
| 34 | sterol metabolic process | 1.92 | 0.00 | 0.40 |
| 35 | regulation of receptor activity | 1.92 | 0.01 | 0.40 |
| 36 | negative regulation of cellular macromolecule biosynthetic process | 1.92 | 0.00 | 0.39 |
| 37 | 2'-deoxyribonucleotide metabolic process | 1.92 | 0.01 | 0.38 |
| 38 | regulation of production of molecular mediator of immune response | 1.91 | 0.00 | 0.39 |
| 39 | positive regulation of histone h3-k4 methylation | 1.90 | 0.00 | 0.40 |
| 40 | steroid metabolic process | 1.90 | 0.01 | 0.39 |

### COPD

Table 12: The 40 highest-scoring gene sets ranked by TEMPO score for the COPD data set

| Rank | Gene Set | Control MSE | Score | MSE p-value | Score p-value | Score BH |
|---|---|---|---|---|---|---|
| 1 | alanine transport | 0.430 | 305.978 | 0.002 | 0.002 | 0.076 |
| 2 | positive regulation of interferon-gamma secretion | 0.145 | 272.491 | 0.002 | 0.004 | 0.093 |
| 3 | positive regulation of phospholipid biosynthetic process | 2.419 | 203.185 | 0.004 | 0.002 | 0.076 |
| 4 | T-helper cell lineage commitment | 5.208 | 194.402 | 0.004 | 0.004 | 0.093 |
| 5 | T-helper 17 cell lineage commitment | 5.208 | 194.402 | 0.004 | 0.006 | 0.093 |
| 6 | transcytosis | 0.759 | 192.616 | 0.002 | 0.002 | 0.076 |
| 7 | vascular smooth muscle cell development | 2.600 | 177.217 | 0.004 | 0.002 | 0.076 |
| 8 | nephric duct development | 1.756 | 175.896 | 0.002 | 0.002 | 0.076 |
| 9 | mRNA transcription from RNA polymerase II promoter | 1.468 | 137.089 | 0.002 | 0.002 | 0.076 |
| 10 | opioid receptor signaling pathway | 7.566 | 128.480 | 0.028 | 0.008 | 0.095 |
| 11 | regulation of extracellular matrix organization | 6.449 | 122.182 | 0.030 | 0.002 | 0.076 |
| 12 | epithelial tube branching involved in lung morphogenesis | 1.066 | 109.133 | 0.002 | 0.008 | 0.095 |
| 13 | RNA surveillance | 4.160 | 106.883 | 0.014 | 0.006 | 0.093 |
| 14 | purine nucleobase transport | 3.093 | 98.951 | 0.004 | 0.010 | 0.095 |
| 15 | monocyte chemotaxis | 4.356 | 96.412 | 0.008 | 0.006 | 0.093 |
| 16 | cell proliferation in forebrain | 6.000 | 95.990 | 0.014 | 0.002 | 0.076 |
| 17 | regulation of cell-cell adhesion mediated by cadherin | 5.771 | 93.285 | 0.012 | 0.006 | 0.093 |
| 18 | negative regulation of extracellular matrix organization | 4.058 | 88.945 | 0.006 | 0.008 | 0.095 |
| 19 | regulation of protein complex stability | 5.739 | 87.307 | 0.010 | 0.008 | 0.095 |
| 20 | regulation of oligodendrocyte differentiation | 4.785 | 86.896 | 0.010 | 0.004 | 0.093 |
| 21 | adrenal gland development | 8.023 | 85.309 | 0.048 | 0.006 | 0.093 |
| 22 | glutamine family amino acid metabolic process | 3.983 | 83.608 | 0.004 | 0.002 | 0.076 |
| 23 | sulfide oxidation | 4.087 | 83.123 | 0.010 | 0.010 | 0.095 |
| 24 | sulfide oxidation, using sulfide:quinone oxidoreductase | 4.087 | 83.123 | 0.004 | 0.012 | 0.105 |
| 25 | regulation of cardiac muscle cell membrane potential | 5.520 | 81.049 | 0.012 | 0.010 | 0.095 |
| 26 | pyrimidine-containing compound transmembrane transport | 2.909 | 79.713 | 0.004 | 0.006 | 0.093 |
| 27 | neg. reg. of mitochondrial outer membrane permeabilization involved in apoptotic signaling pathway | 4.213 | 79.237 | 0.008 | 0.010 | 0.095 |
| 28 | negative regulation of activin receptor signaling pathway | 6.134 | 78.457 | 0.018 | 0.014 | 0.105 |
| 29 | regulation of microvillus organization | 5.683 | 77.186 | 0.014 | 0.014 | 0.105 |
| 30 | regulation of microvillus assembly | 5.683 | 77.186 | 0.014 | 0.012 | 0.105 |
| 31 | glutamine transport | 3.735 | 77.005 | 0.014 | 0.026 | 0.106 |
| 32 | neuroblast proliferation | 7.298 | 75.503 | 0.038 | 0.006 | 0.093 |
| 33 | sequestering of metal ion | 5.122 | 73.510 | 0.026 | 0.026 | 0.106 |
| 34 | negative regulation of protein sumoylation | 3.962 | 73.403 | 0.004 | 0.020 | 0.106 |
| 35 | nucleobase-containing small molecule interconversion | 1.875 | 71.447 | 0.002 | 0.006 | 0.093 |
| 36 | bundle of His cell-Purkinje myocyte adhesion involved in cell communication | 10.007 | 68.231 | 0.040 | 0.016 | 0.106 |
| 37 | positive regulation of Rho protein signal transduction | 5.440 | 62.829 | 0.018 | 0.010 | 0.095 |
| 38 | response to acid chemical | 2.122 | 58.860 | 0.002 | 0.002 | 0.076 |
| 39 | reg. of cardiac muscle contraction by calcium ion signaling | 3.821 | 58.222 | 0.004 | 0.004 | 0.093 |
| 40 | bicellular tight junction assembly | 7.333 | 57.457 | 0.044 | 0.006 | 0.093 |

Table 13: The 40 highest-scoring upregulated gene sets returned by GSEA in COPD.

| Rank | Gene Set | NES | p-value | FDR |
|---|---|---|---|---|
| 1 | peptidyl-proline hydroxylation | -1.917 | 0.002 | 1.000 |
| 2 | positive regulation of nuclease activity | -1.838 | 0.002 | 1.000 |
| 3 | regulation of isotype switching | -1.834 | 0.000 | 1.000 |
| 4 | peptidyl-serine autophosphorylation | -1.827 | 0.000 | 1.000 |
| 5 | oocyte differentiation | -1.812 | 0.004 | 1.000 |
| 6 | sulfide oxidation | -1.761 | 0.002 | 1.000 |
| 7 | sulfide oxidation, using sulfide:quinone oxidoreductase | -1.761 | 0.002 | 1.000 |
| 8 | regulation of mitophagy | -1.750 | 0.000 | 1.000 |
| 9 | macromolecule depalmitoylation | -1.746 | 0.002 | 1.000 |
| 10 | iron-sulfur cluster assembly | -1.745 | 0.000 | 1.000 |
| 11 | metallo-sulfur cluster assembly | -1.745 | 0.000 | 1.000 |
| 12 | protein hydroxylation | -1.732 | 0.000 | 1.000 |
| 13 | regulation of torc1 signaling | -1.729 | 0.006 | 1.000 |
| 14 | response to misfolded protein | -1.716 | 0.004 | 1.000 |
| 15 | negative regulation of macroautophagy | -1.711 | 0.002 | 1.000 |
| 16 | mitochondrial atp synthesis coupled proton transport | -1.704 | 0.046 | 1.000 |
| 17 | positive regulation of ruffle assembly | -1.697 | 0.018 | 1.000 |
| 18 | oocyte development | -1.696 | 0.010 | 1.000 |
| 19 | positive regulation of isotype switching | -1.695 | 0.008 | 1.000 |
| 20 | activation of signaling protein activity involved in unfolded protein response | -1.686 | 0.011 | 1.000 |
| 21 | nucleosome disassembly | -1.685 | 0.002 | 1.000 |
| 22 | chromatin disassembly | -1.685 | 0.002 | 1.000 |
| 23 | protein-dna complex disassembly | -1.685 | 0.002 | 1.000 |
| 24 | positive regulation of erad pathway | -1.680 | 0.000 | 1.000 |
| 25 | energy coupled proton transport, down electrochemical gradient | -1.679 | 0.048 | 1.000 |
| 26 | atp synthesis coupled proton transport | -1.679 | 0.048 | 1.000 |
| 27 | base-excision repair, ap site formation | -1.677 | 0.011 | 1.000 |
| 28 | peptidyl-diphthamide metabolic process | -1.676 | 0.011 | 1.000 |
| 29 | peptidyl-diphthamide biosynthetic process from peptidyl-histidine | -1.676 | 0.011 | 1.000 |
| 30 | positive regulation of neurotransmitter secretion | -1.672 | 0.004 | 1.000 |
| 31 | mitochondrial respiratory chain complex assembly | -1.670 | 0.037 | 1.000 |
| 32 | positive regulation of anoikis | -1.669 | 0.006 | 1.000 |
| 33 | establishment of protein localization to mitochondrion | -1.668 | 0.034 | 1.000 |
| 34 | regulation of autophagosome maturation | -1.663 | 0.004 | 1.000 |
| 35 | protein localization to mitochondrion | -1.661 | 0.026 | 1.000 |
| 36 | regulation of immunoglobulin production | -1.657 | 0.012 | 1.000 |
| 37 | histone h4-k12 acetylation | -1.648 | 0.002 | 1.000 |
| 38 | calcineurin-nfat signaling cascade | -1.643 | 0.011 | 1.000 |
| 39 | regulation of nuclease activity | -1.643 | 0.006 | 1.000 |
| 40 | heme biosynthetic process | -1.639 | 0.015 | 1.000 |

Table 14: The 40 highest-scoring upregulated gene sets returned by maSigPro+GSEA preranked in COPD.

| Rank | Gene Set | NES | p-value | FDR |
|---|---|---|---|---|
| 1 | ectodermal placode development | 2.104 | 0.000 | 0.091 |
| 2 | ectodermal placode morphogenesis | 2.102 | 0.000 | 0.047 |
| 3 | ectodermal placode formation | 2.077 | 0.000 | 0.049 |
| 4 | response to amino acid | 1.988 | 0.000 | 0.117 |
| 5 | peptidyl-methionine modification | 1.966 | 0.001 | 0.124 |
| 6 | positive regulation of er-associated ubiquitin-dependent protein catabolic process | 1.934 | 0.000 | 0.147 |
| 7 | ubiquinone biosynthetic process | 1.933 | 0.000 | 0.130 |
| 8 | base-excision repair, ap site formation | 1.930 | 0.000 | 0.117 |
| 9 | positive regulation of intracellular estrogen receptor signaling pathway | 1.922 | 0.001 | 0.112 |
| 10 | cellular response to amino acid stimulus | 1.917 | 0.000 | 0.109 |
| 11 | spliceosomal tri-snrnp complex assembly | 1.914 | 0.000 | 0.103 |
| 12 | protein monoubiquitination | 1.906 | 0.000 | 0.105 |
| 13 | histone ubiquitination | 1.903 | 0.000 | 0.099 |
| 14 | quinone biosynthetic process | 1.891 | 0.000 | 0.106 |
| 15 | ubiquinone metabolic process | 1.887 | 0.000 | 0.104 |
| 16 | short-chain fatty acid catabolic process | 1.883 | 0.001 | 0.101 |
| 17 | jun phosphorylation | 1.881 | 0.002 | 0.099 |
| 18 | acetyl-coa biosynthetic process from pyruvate | 1.871 | 0.000 | 0.105 |
| 19 | regulation of establishment of protein localization to telomere | 1.869 | 0.003 | 0.102 |
| 20 | trna threonylcarbamoyladenosine metabolic process | 1.864 | 0.001 | 0.101 |
| 21 | mitochondrial rna modification | 1.862 | 0.002 | 0.100 |
| 22 | intermediate filament organization | 1.861 | 0.001 | 0.096 |
| 23 | regulation of establishment of protein localization to chromosome | 1.852 | 0.001 | 0.103 |
| 24 | histone monoubiquitination | 1.849 | 0.000 | 0.102 |
| 25 | smad protein complex assembly | 1.843 | 0.000 | 0.104 |
| 26 | regulation of autophagosome maturation | 1.840 | 0.001 | 0.104 |
| 27 | spliceosomal snrnp assembly | 1.839 | 0.000 | 0.101 |
| 28 | mitochondrial rna processing | 1.839 | 0.000 | 0.098 |
| 29 | mitochondrial trna modification | 1.836 | 0.002 | 0.097 |
| 30 | depyrimidination | 1.832 | 0.000 | 0.098 |
| 31 | positive regulation of protein localization to cajal body | 1.819 | 0.000 | 0.108 |
| 32 | regulation of mrna 3'-end processing | 1.816 | 0.000 | 0.108 |
| 33 | regulation of low-density lipoprotein particle clearance | 1.812 | 0.003 | 0.109 |
| 34 | regulation of protein localization to cajal body | 1.808 | 0.001 | 0.111 |
| 35 | pyrimidine-containing compound transmembrane transport | 1.797 | 0.001 | 0.121 |
| 36 | mitochondrion morphogenesis | 1.797 | 0.000 | 0.118 |
| 37 | nucleobase-containing small molecule catabolic process | 1.797 | 0.002 | 0.115 |
| 38 | primary mirna processing | 1.795 | 0.001 | 0.114 |
| 39 | membrane depolarization during sa node cell action potential | 1.792 | 0.009 | 0.115 |
| 40 | viral release from host cell | 1.791 | 0.000 | 0.114 |
